# Conserved Rhizosphere Microbiomes and Metabolic Functions Across Diverse Grapevine Rootstocks: Implications for Plant Elemental Composition

**DOI:** 10.64898/2026.01.14.698671

**Authors:** Joel F. Swift, Grace E. Trello, Zachary N. Harris, Zoë Migicovsky, Allison J. Miller

## Abstract

**Background:** While grapevine rootstocks are essential to commercial viticulture, researchers still lack a complete understanding of how these root systems and their associated microbiomes influence the vine’s mineral nutrition. Previous research has shown that rootstocks with similar pedigrees, reflecting the limited number of parental genotypes in the breeding pool, affect both the vine’s elemental profile and microbiome composition. Here, we extend this work by surveying a broader diversity of rootstock genotypes representing distinct parentage groups across multiple scions and by assessing both functional and taxonomic associations between the rhizosphere metagenome and vine elemental composition.

**Results:** We used a rootstock diversity trial with ten rootstock genotypes grafted to one of two scions to simultaneously characterize the rhizosphere metagenome and elemental composition of both berries and roots. We found that elemental composition varied strongly among vine compartments, while scion and rootstock had comparatively minor effects. Berries of ‘Cabernet Sauvignon’ contained higher concentrations of boron, potassium, and phosphorus than those of ‘Chardonnay’. Only three elements in root tissues differed among rootstock genotypes, and these differences were unrelated to genetic relatedness among rootstocks. Rhizosphere microbiome composition was also conserved across rootstocks, suggesting that diverse genotypes recruit a consistent microbial community with similar functional potential. By linking rhizosphere microbiome profiles with root elemental composition, we found that *Streptomyces* and *Mesorhizobium* were significantly associated with elemental patterns, showing negative correlations with a principal component heavily loaded for molybdenum, cadmium, potassium, and iron.

**Conclusions:** Our findings suggest that vine compartment exerts a dominant influence on elemental composition, while rootstock genotypes have comparatively modest effects. Despite the overall conservation of the rhizosphere microbiome, specific microbial taxa were linked to elemental variation, highlighting potential microbe-element associations that may influence nutrient dynamics in the grapevine rhizosphere. These results provide a foundation for disentangling how root-associated microbes contribute to vine mineral nutrition, or vice versa, across diverse rootstock genotypes.

## Introduction

Grafting in viticulture involves the union of scion (shoot system) and rootstock (root system) to form a single plant. Over 80% of commercially grown grapevines in the world are grafted (Ollat *et al*., 2016) and yet researchers still lack a complete understanding of how rootstocks and their associated microbiomes affect mineral nutrition. Grafting, a centuries-old horticultural practice adopted to combat *Phylloxera* (Mudge *et al*., 2009), continues to serve as a powerful tool for improving vine performance. As breeding new grapevine varieties is time-consuming and costly, rootstocks provide an accessible and speedy means of modifying vine phenotypes. Currently, grafting and rootstock selection are utilized globally to manage vigor, alter fruit quality, and improve tolerance to abiotic stress (Warschefsky *et al*., 2016). More recently, grafting has been demonstrated to influence a host of multidimensional phenotypes including the metabolome (Zhong *et al*., 2022), gene expression (Harris *et al*., 2023), the microbiome (Swift *et al*., 2021, 2023), and elemental composition (Harris *et al*., 2022).

Roots are the primary organ responsible for the acquisition of most nutrients. Many studies profiling the elemental composition of grapevines have found elemental profiles are influenced by both the rootstock (Bavaresco *et al*., 2003; Lecourt *et al*., 2015; Migicovsky *et al*., 2019; Gautier *et al*., 2020; Harris *et al*., 2021) and scion genotypes (Verdugo-Vásquez *et al*., 2021). This has led to discoveries such as ‘Chambourcin’ vines grafted to the rootstock cultivar ‘SO4’ accumulating higher foliar levels of molybdenum and nickel compared to other rootstocks (Harris *et al*., 2021) and rootstock influence on nitrogen, phosphorus, and potassium across multiple scions (Ibacache & Sierra, 2009). This modulation of scion elemental composition is likely a product of complex rootstock genotype x environment interactions. For example, because most major rootstocks are derived from the same three wild *Vitis* species (*Vitis riparia* (Michx.), *V. rupestris* (Scheele), and *V. berlandieri* (Planch.)), it is possible to determine whether rootstock parentage (a proxy for genetic effects) influences elemental composition. Higher concentrations of phosphorus at anthesis in the scion have been associated with *V. berlandieri* and *V. rupestris*, and *V. riparia* parentage is associated with increased concentration of sulfur and decreased concentrations of phosphorus and magnesium in leaves of the grafted scion (Gautier *et al*., 2020), to the point of deficiency with some rootstock genotypes (Morel *et al*., 2024). Moreover, rootstock genotype interacted with temporal and environmental covariates to influence concentrations of 20 elements surveyed in scion across three years (Harris *et al*., 2022). While there are clear connections between rootstock genotype, environment, and foliar elemental composition, open questions remain about the mechanisms underlying these associations and the elemental composition of non-leaf tissue. The root-associated microbiome represents a plausible intermediary, potentially mediating the availability and uptake of mineral nutrients.

Although compartment specificity is a strong differentiator, both rootstock and scion genotypes contribute to variation in microbial community composition across grapevine compartments (Gautier *et al*., 2020; Vink *et al*., 2021; Marasco *et al*., 2022; Swift *et al*., 2023). Graft combinations can influence microbial community structure and may also shape the functional processes mediated by these microbes, including nutrient cycling in the rhizosphere. Microorganisms and their genomes, collectively termed the metagenome (Riesenfeld *et al*., 2004), harbor diverse metabolic pathways for converting elements, notably nitrogen, phosphorus, and sulfur, to forms that are available for uptake by plants (Bonkowski, 2004; Rodríguez *et al*., 2006; Richardson *et al*., 2009; Jacoby *et al*., 2017; Dai *et al*., 2020). Thus, much of the research on the metagenomics of rhizospheres has focused on identifying the functional connections between the microbiome and plant nutrition (Salas-González *et al*., 2021). For example, iron, which serves as an important co-factor in many enzymes, is required for the synthesis of chlorophyll (Rout & Sahoo, 2015; Schmidt *et al*., 2020) and plays a role in plant-microbe interactions. Many plant species show enrichment of *Actinobacteria* in the rhizosphere when under drought stress (Santos-Medellín *et al*., 2017; Naylor *et al*., 2017; Naylor & Coleman-Derr, 2018; Xu *et al*., 2018a; Santos-Medellín *et al*., 2021); recently, this phenomenon was linked to plant-driven decreases in iron availability and abiotic changes in the soil (Xu *et al*., 2021). As drought intensified, genes involved in iron uptake in sorghum were downregulated. Combined with the more aerobic soil conditions under drought, this reduced the availability of soluble iron in the rhizosphere, favoring *Actinobacteria* spp., which outcompete other taxa under iron limitation. Additional roles have been uncovered for rhizosphere associated microorganisms in plant elemental concentrations, including zinc (Krithika & Balachandar, 2016; Singh *et al*., 2018; Wang *et al*., 2021), iron (Jia *et al*., 2022), and phosphorus (Unno & Shinano, 2013). Thus, combining both metagenomics and elemental profiling provides a powerful tool for understanding the complex nature of the plant rhizosphere.

To explore plant-microbe interactions and their influence on elemental concentrations in roots and shoots, we used a diverse set of rootstock genotypes in an experimental vineyard in San Joaquin County, California. This vineyard included 10 rootstock genotypes each grafted to ‘Cabernet Sauvignon’ and ‘Chardonnay’ scions (Table 1). We applied shotgun metagenomics to assess the taxonomic and functional capacity of rhizosphere microbiomes and elemental analyses to identify links between the root microbiome and vine elemental composition. Specifically, we sought to answer the following questions: **1)** What are the major factors shaping elemental composition across berry and root tissues? **2)** Do rootstock and/or scion cultivars assemble distinct rhizosphere microbiomes and are there conserved taxa and functions across cultivars? And **3)** Do elemental profiles for the rhizosphere correlate with changes in the microbiome composition or functional capacity?

## Materials and Methods

### Experimental Design and Sampling

Samples were collected from a rootstock diversity trial vineyard in Central California (San Joaquin County). Vines were cultivated with a simple curtain trellis design setup in a north-south orientation, with vines and rows spaced 2.1 m and 3 m apart, respectively. The soil texture varied from loam to sandy loam. The vineyard was established in 1994 and featured 10 rootstock cultivars (Table 1) grafted to either ‘Cabernet Sauvignon’ or ‘Chardonnay’. The vineyard used a split-block randomized design (*i.e.*, two blocks, one per scion), with constant water levels maintained throughout. Each scion/rootstock combination was replicated at least four times per block, with each replicate containing eight or nine vines. Additional vineyard establishment and management details are provided in Migicovsky *et al*. (2021).

From each scion/rootstock combination, we collected root and berry samples from three replicate vines (*n* = 60 per compartment; 10 *rootstocks* × 2 *scions* × 3 *replicate vines/rootstock-scion combination*). Root samples were carefully excavated using an ethanol-sterilized trowel (95% v/v) at a depth of 20-30 cm, removed by hand utilizing sterilized gloves. Loose soil particles were removed by shaking. Berries were collected as an intact cluster, clipping the rachis of the cluster at the point of attachment with the shoot using sterilized stainless-steel pruning shears, and stored in a plastic bag. Samples were placed into coolers with ice packs immediately in the field for transport to the lab. Our sampling strategy sought to capitalize on the vineyard design by maximizing the distance between replicates, thus minimizing the impact of intra-vineyard heterogeneity in soil microbiota or properties. The majority of the collections (n = 135) were made on July 23rd, 2018, with a smaller number (n = 19) recollected on July 23rd, 2019 due to damage during shipment from California to Missouri. Two individual berry samples were unrecoverable, ‘Chardonnay’ grafted to ‘3309 Couderc’ and ‘Cabernet Sauvignon’ grafted to ‘101-14 MGT’. Samples were stored at -80 °C until processing.

### Rhizosphere elemental composition

A portion of each root sample (with rhizosphere intact) and several berries (from cardinal directions on the cluster: top, tip, opposite sides) were dried to a constant weight at 70 °C. Individual samples were crushed to a fine powder with a mortar and pestle. 75-100 mg of each sample was used to quantify ion concentrations. Briefly, samples were acid digested and subjected to inductively coupled plasma mass spectrometry. All procedures followed the protocol of the

Donald Danforth Plant Science Center Ionomics Pipeline (Baxter, 2010; Ziegler *et al*., 2013). Ion concentrations were corrected for initial sample mass and instrument drift, using internal standards. The final dataset provided concentrations for 19 ions: aluminum, arsenic, boron, calcium, cadmium, cobalt, copper, iron, potassium, magnesium, manganese, molybdenum, sodium, nickel, phosphorus, rubidium, selenium, strontium, and zinc. Due to analytical issues, concentrations for sulfur (typically considered in the established pipeline) were not quantified.

### Rhizosphere extraction and microbiome sequencing

For rhizosphere extractions, ∼45 mL of freshly prepared and sterile phosphate buffered saline (PBS; amended with 0.001% Silwet-77 v/v) was added to the 50 mL tube containing roots and vortexed for 2 minutes and agitated for 30 minutes at 180 rpm on a shaking platform. Root tissue was removed from tubes, and the samples were vortexed briefly. The slurry was subjected to centrifugation at 4,000 g for 10 minutes. The supernatant (PBS) was decanted, and the resulting rhizosphere pellet was transferred to a 2 mL tube, which was centrifuged at 21,000 g for 3 minutes. The pellet was frozen at –80 °C after removing residual supernatant.

DNA extractions were performed at the Danforth Plant Science Center (Saint Louis, MO, USA) using the DNeasy PowerSoil Pro Kit (Qiagen, Germantown, MD, USA) following the manufacturer’s protocol with slight modification: we used a starting sample weight of 350 mg and a 30-minute incubation at 65 °C incubation prior to homogenization. Samples were further cleaned using a DNeasy PowerClean Pro Clean Up Kit according to the manufacturer’s protocol. All extracts were quantified using a DS-11 spectrophotometer (Denovix, Wilmington, DE, USA). Shotgun metagenomic libraries were prepared by Novogene (Sacramento, CA) and sequencing was conducted on a NovaSeq 6000 (Illumina, San Diego, CA) with a 2 x 150bp flowcell.

### Sequence Processing

Sequence files were examined for quality using FastQC (www.bioinformatics.babraham.ac.uk/projects/fastqc/) and the resulting outputs were concatenated into a single HTML report utilizing MultiQC (Ewels *et al*., 2016). We utilized the bioBakery “Whole Metagenome Shotgun” workflow (Mclver *et al*., 2018; Beghini *et al*., 2021). In brief, KneadData v0.7.4 was used to perform quality control, sequence files were trimmed of adapters and a sliding window approach was utilized to remove low-quality bases at the ends of reads using Trimmomatic (Bolger *et al*., 2014). Sequence files were then decontaminated by mapping reads to the *Vitis vinifera* cv. PN40024 12X.v2 reference genome (Canaguier *et al*., 2017) using Bowtie2 (Langmead & Salzberg, 2012); any reads with significant alignments to the reference genome were discarded. After quality control and decontamination, the taxonomic profile of each sample was assessed via MetaPhlAn v3.0.13 and the functional profile (gene families and metabolic pathways) was assessed via HUMAnN v3.0.0 (Beghini *et al*., 2021). Both programs were run with default settings except for the ‘stat_q’ setting in MetaPhlAn which was lowered to 0.1 to increase the number of markers considered when attempting taxonomic classification and relative abundance calculation. Taxonomic and functional profiles were normalized for sequencing depth (total sum scaling) via the ‘humann_renorm_table’ function. Taxonomic profiles were converted to relative abundance, and functional profiles to copies per million (CPM), which accounts for gene length variation and number of mapped reads, for downstream analyses.

### Elemental Analysis

All statistical analysis was conducted in R v4.2.2 (R Core Team, 2025), utilizing the following packages: tidyverse v2.0.0 (Wickham *et al*., 2019), lme4 v1.1-351 (Bates *et al*., 2015), emmeans v1.9.0 (Lenth *et al*., 2020), corrplot v0.92 (Wei & Simko, 2021), ggpubr v0.6.0 (Kassambara, 2023), and factoextra v1.0.7 (Kassambara & Mundt, 2020). Samples with elemental concentrations greater than five standard deviations from the mean were removed prior to the analysis. Due to the need to recollect samples, we first sought to assess whether the year the samples were collected influenced the elemental composition. Elemental concentrations were z-score transformed and these values were the basis for linear models. Each element was modeled with the following model (Eq1):

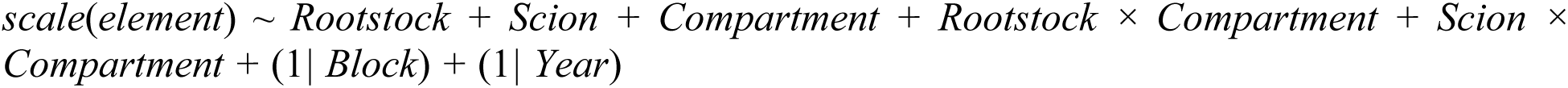

In this model, the “block” term represents the four independent plots per scion block, serving as a proxy for vineyard location. We found that the random effect of sampling year was non-significant for each elemental model (Likelihood Ratio Test) and no clustering of samples by year in principal component space (Figure S1, Table S2). Thus, we choose to include samples from 2018 and 2019 in all analyses and model collection year as a random effect in all models.

For each element, we fit the model (above) to its scaled concentration. P-values were computed for each term under a type-3 ANOVA framework, with P-value correction where appropriate using false discovery rate (Benjamini Hochberg) correction. Model assumptions of normality were assessed by inspecting model residuals (Harrison *et al*., 2018). For significant model terms and interactions, post hoc comparisons of estimated marginal means were tested using emmeans with a Tukey-correction for multiple comparisons. Principal component analysis (PCA) was used to visualize the potential groupings caused by experimental factors. Biplots were generated to inspect the loadings of elements. A Pearson’s correlation matrix per compartment was generated to explore elemental correlations.

As the rootstocks we studied are of hybrid origin, we elected to consider rootstocks first as categorical (*i.e*., each rootstock as its own group), we then modeled rootstocks in groups by parentage (Table S1), and finally by the genetic distance between rootstock pairs. For parentage-based analyses, we grouped rootstocks into 3 groups; *V. riparia* × *V. rupestris* (n = 3), *V. berlandieri* × *V. rupestris* (n = 4), *V. berlandieri* × *V. riparia* (n = 3). For genetic distance-based analyses, we utilized 21 nuclear microsatellite markers from Riaz *et al*. (2019). This study sought to clarify parentage assignments across 36 viticulturally important rootstocks. The microsatellite table was imported into R, converted to a “genind” object (genotypes of individuals), and PopGenReport v3.1 (Adamack & Gruber, 2014) was used to calculate a genetic distance matrix. We elected to utilize Kosman diversity, which allows for missing data, to measure genetic distances between individual rootstocks (Kosman & Leonard, 2005). A Euclidean distance matrix was constructed for each plant compartment with element concentrations averaged for a given rootstock. A Mantel test assessed the correlation between the genetic distance matrix and elemental composition Euclidean distance matrix.

### Microbiome Analysis

For taxonomic profiles, we used principal coordinate analysis (PCoA) with Bray-Curtis dissimilarity calculated from the package phyloseq v1.36.0 (McMurdie & Holmes, 2013) in order to visualize clustering of samples across experimental factors. PERMANOVA analyses were conducted using the package vegan v2.5.7 (Oksanen *et al*., 2019) via the function ‘Adonis’ and fit using a model with rootstock, scion, and their interaction as marginal fixed effects and Bray-Curtis dissimilarity as the response with 1,000 permutations. We tested for differential taxonomic abundance using MaAsLin2 v1.6.0 (Mallick *et al*., 2021). Each taxonomic level was tested with the same model (Eq2):

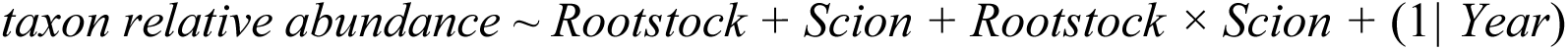

P-values were corrected for multiple tests using false discovery rate (Benjamini Hochberg) correction. The functional capacity of the rhizospheres of rootstock/scion combinations was also assessed via MaAsLin2, which fit the same model to each pathway (Eq3):

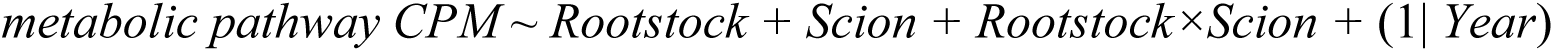

### Defining core taxa and functional capacities of the rhizosphere microbiome

To define core members of the microbiome and core functions of the microbiome, we examined both the occupancy and relative abundance of each feature (either taxa or pathway). As the filtering parameters for defining a core microbiome from the literature vary widely (Shade & Stopnisek, 2019), we choose filtering parameters in line with Walters *et al*., (2018) that were conservative in order to reduce noise. For defining core taxa, we selected genera that were present in ≥95% of samples (i.e., present in ≥57 of 60 samples) and had a mean relative abundance of ≥0.4% (80^th^ percentile). For defining core functional capacities, we selected pathways that were present in 100% of samples and had a mean count greater than 4215 CPM (80^th^ percentile). We chose to increase the occupancy requirement for defining core functional capacities as we observed a larger number of pathways were conserved across all samples compared to taxa.

### Associating elemental profiles with the rhizosphere microbiome

We found a strong correlation structure among elemental concentrations and thus opted to utilize principal components (PCs) to test for associations with the rhizosphere microbiome. A linear model was fit to the first 10 principal components (cumulatively capturing 94.3% of the variance) using the following model (Eq4):

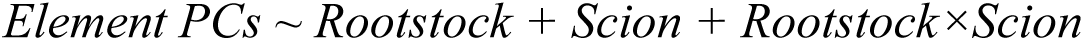

We utilized elemental PCs to test for associations with both the taxa and metabolic pathways. For taxa, we used genus-level assignments as a balance between over- and under-splitting. We filtered both genera and pathways to an occupancy of ≥50% of the samples. For genera, we additionally required that each genus had a mean relative abundance >0.5%. We used MaAsLin2 to fit a model with the elemental PCs each as fixed effects and year included as a random effect. Scatterplots were created for genera and pathways that were significantly correlated with elemental PCs.

## Results

### Elemental concentrations vary by compartment, with minor effects of scion and rootstock

Root and berry compartments show distinct elemental compositions. Across principal component 1 (68.4%), samples clearly clustered by compartment (Figure 2A). Correspondingly, all elements except phosphorus varied significantly across compartments (Figure S2; Table S3) and compartment was the largest explainer of variance for all elements (5.9 – 82.8% variance explained). Berries contained greater concentrations of boron (9.78 vs 3.47 ppm) and potassium (15393 vs 4447 ppm) compared to roots from the same vine (Figure S2). Conversely, roots contained greater concentrations of multiple macro- and micro-nutrients (e.g., calcium, magnesium, iron, and manganese, among others) and heavy metals (Figure S2). Elemental profiles also showed differing correlations between compartments (Figure 2B). Within root samples, many heavy metals were negatively correlated with the macronutrient phosphorus. In contrast, berries had no significant negative correlations between elements, with the highest correlations between iron and aluminum (r = 0.97) and calcium and strontium (0.95). Comparing the root and berry elemental concentrations, on an element-by-element basis within individual vines, we found a moderate correlation overall (r = 0.56 ± 0.15), but permutation testing showed this to be non-significant (*p* = 0.79). Overall, this suggests compartments’ elemental profiles are strongly regulated by the plant, with berries and roots showing different allocation and correlation patterning.

**Figure 1.**
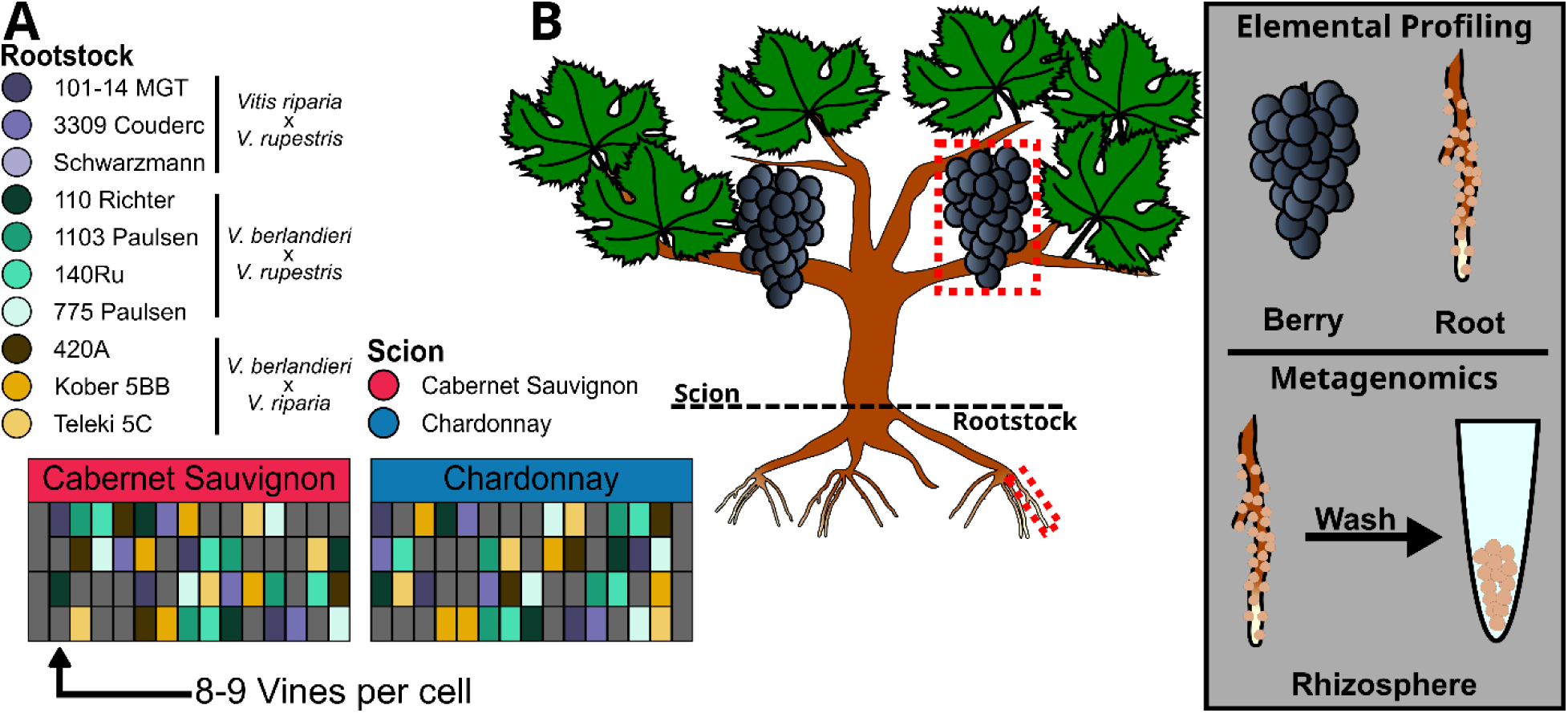
Experimental design and sample collection per vine. **A)** Samples were collected from a rootstock diversity trial vineyard near Lodi in San Joaquin County, California. Ten rootstock genotypes were selected from the vineyard for collection; each is a cross between two of three *Vitis spp.*; *V. berlandieri*, *V. riparia*, and *V. rupestris*^†^. Each rootstock was grafted to either a ‘Chardonnay’ (selection FPS 04) or ‘Cabernet Sauvignon’ (selection FPS 07) scion. A vineyard map depicts the split-block design. Colored cells denote rootstock genotypes, and gray cells represent vines not utilized in the study. **B)** From each vine, we collected an intact berry cluster and several root segments with rhizosphere attached. These collections were used as the basis to conduct elemental profiling for berries and roots, and metagenomic sequencing for the rhizosphere. ^†^Additional parentage information for rootstock genotypes is provided in Table S1.

**Figure 2.**
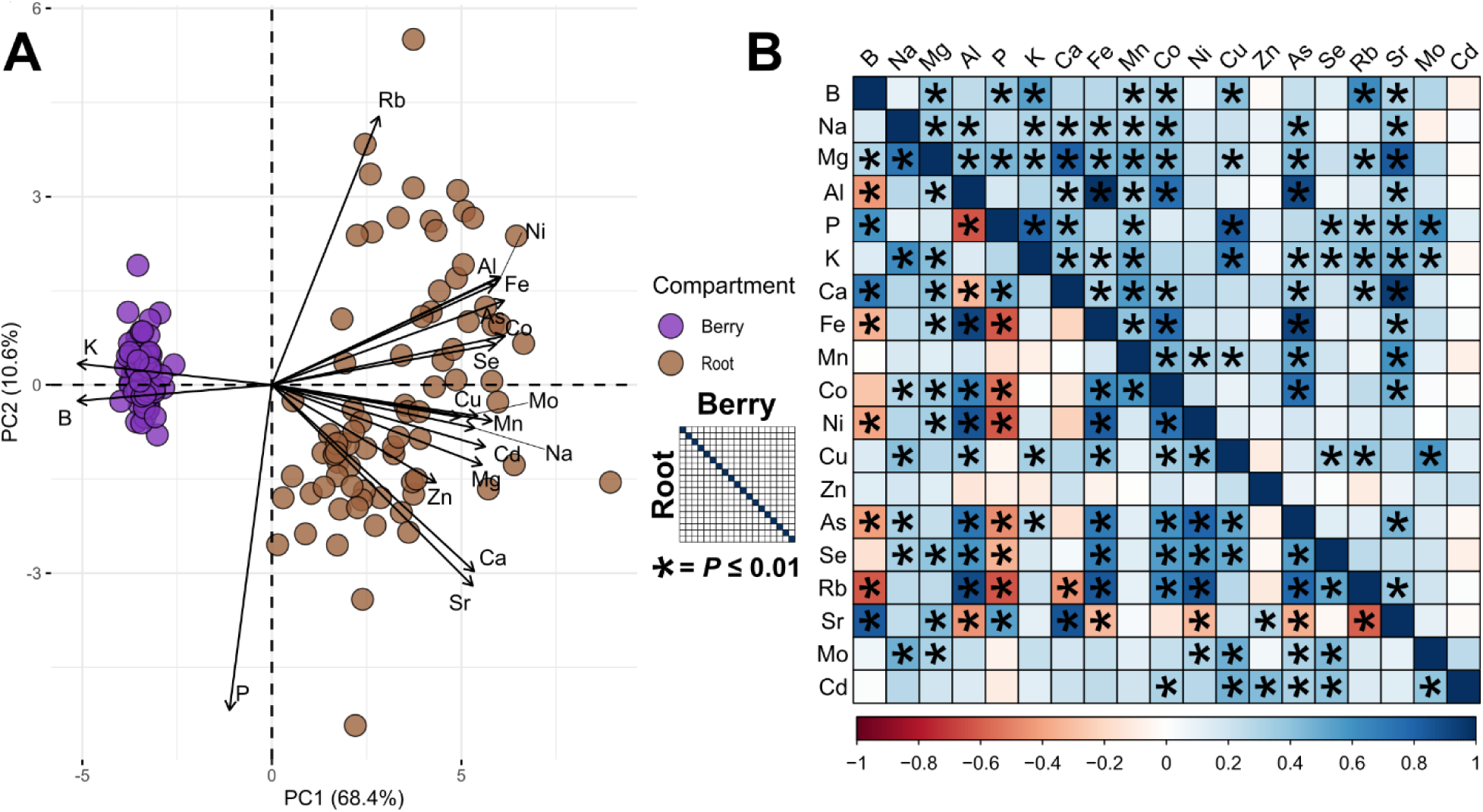
Elemental profiles are strongly differentiated by compartment. **A)** Principal component analysis with arrows representing loadings of elements on principal components 1 and 2. **B)** Pearson’s correlations among elements for berry (upper diagonal) and root (lower diagonal) samples. Asterisks denote significant correlations (P ≤ 0.01).

Berry elemental profiles were further shaped by the scion cultivar (Figure S3). Visualizing samples in principal component space, scion cultivars diverge across the third principal component (6.0% of the variation; Figure 3A). Examining the loadings of elements on the third principal component revealed three elements that loaded strongly: potassium, phosphorus, and boron (Figure 3B-D). For each of these three elements, concentrations were greater in ‘Cabernet Sauvignon’ compared to ‘Chardonnay’; however, phosphorus did not significantly differ in berries between the cultivars (Figure 3D).

**Figure 3.**
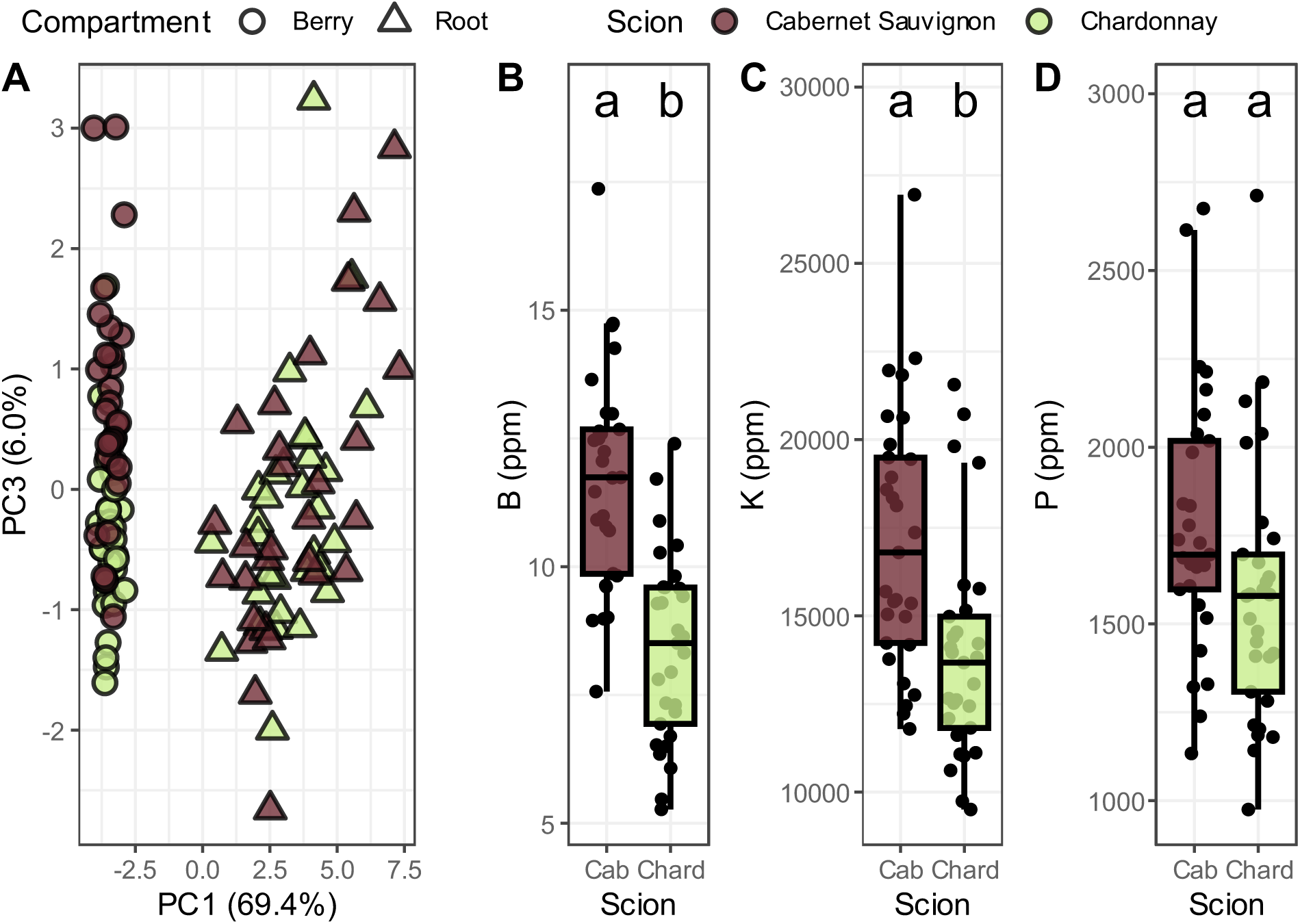
Berry elemental profiles are distinguishable by scion cultivar. **A)** Principal component analysis displaying berry and root samples, point colors represent scion cultivars and shapes represent different plant compartments (n per Compartment-Scion combination = 29-30). Elements that loaded strongly on principal component three were modeled. Berry concentration values for **B)** Boron (ANOVA; *“Compartment×Scion” p* < 0.01, F_1,93_ = 33.49), **C)** Potassium (ANOVA; *“Compartment×Scion” p* = 0.01, F_1,93_ = 6.72), and **D)** Phosphorus (ANOVA; *“Compartment×Scion” p* = 0.44, F_1,93_ = 0.59). Boxplot hinges represent the 1st and 3rd quartiles; whiskers represent 1.5 times the interquartile range; significance letters represent post-hoc comparisons of estimated marginal means.

In contrast to scion cultivar, rootstock genotype had a smaller effect on elemental profiles for either compartment, regardless of whether rootstock was modelled by genotype or by parentage. When treating each rootstock as an independent genotype, three of the element concentrations of roots were impacted by rootstock (manganese, sodium, and phosphorus; Table S3; Figure S4). When rootstocks were grouped by parentage (Table S1), three elements differed in roots by the parentage group (Table S4). The parentage group ‘*V. berlandieri x V. rupestris*’ showed the lowest concentration of all three of the elements (cobalt, manganese, and molybdenum; Figure 4A-C). Each of these elements was generally relegated to the root compartment of the plant (Figure S2). Finally, genetic distance between rootstock genotypes was unrelated to the dissimilarity between elemental profiles of the rootstocks for both compartments (Figure 4D-E). Overall, these results indicate a similar pedigree (*i.e.,* sharing both parental species) was not indicative of a particular elemental profile.

**Figure 4.**
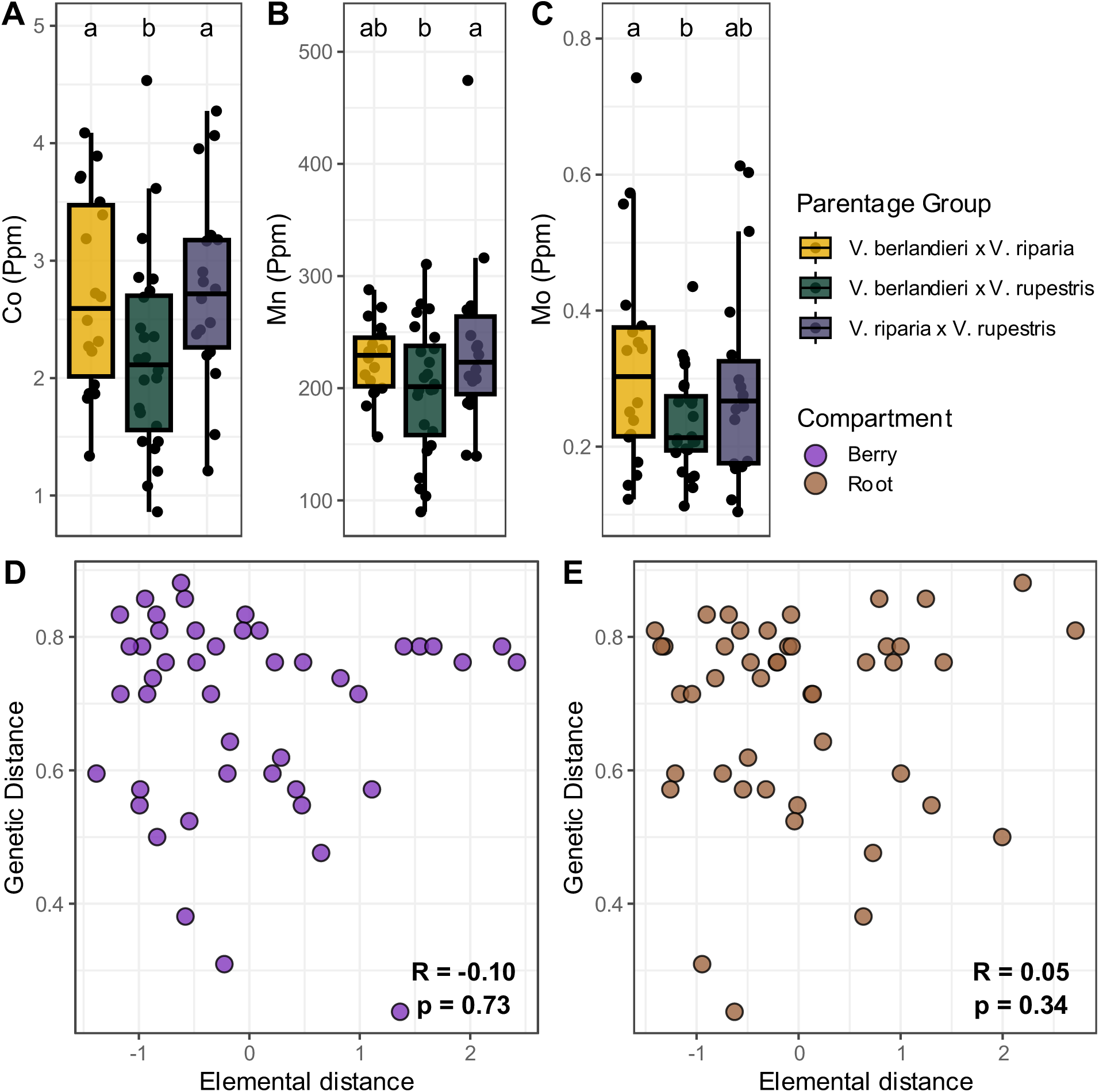
Rootstock cultivar influences elemental composition for specific elements, whereas genetic distance between cultivars does not correspond to differences in elemental profiles. The effect of rootstock parentage group on root concentrations of A) Cobalt (ANOVA; *“Parentage×Compartment” p* = 0.06, F_2,110_ = 2.85), B) Manganese (ANOVA; *“Parentage×Compartment” p* = 0.13, F_2,110_ = 2.06), and C) Molybdenum (ANOVA; *“Parentage×Compartment” p* = 0.4, F_2,110_ = 3.33). Boxplots display elemental concentrations of the root compartment as these elements were in minimal concentrations in berries (see figure S2). Boxplot hinges represent the 1st and 3rd quartiles; whiskers represent 1.5 times the interquartile range; significance letters represent post-hoc comparisons of estimated marginal means. A Mantel test was used to assess the correlation between Euclidean distance of the rootstock’s Z-score transformed elemental profiles (x-axis) to the rootstock’s genetic distance (y-axis). Tests were performed separately for **D)** berry elemental profiles and **E)** root elemental profiles. Permutation tests with 1000 simulations were used to obtain p-values.

### Rhizosphere microbiomes are conserved across rootstock and scion genotypes

Samples from our metagenomic libraries produced a total of 2.37 billion reads with samples ranging from 36.0 to 49.7 million reads (39.5 million per sample mean). After quality filtering, merging of paired reads, and decontamination, we recovered 2.18 billion reads; on average, 6.43% of reads were removed during filtering and an additional 1.61% during decontamination per sample. Although taxonomic profiles showed some variation across rootstock and scion genotypes (Figure 5A–B), PERMANOVA on Bray-Curtis dissimilarity indicated no significant effect of rootstock or scion on the overall rhizosphere community composition (Table S5). We identified prokaryotic members of 13 phyla, 29 classes, 61 orders, 104 families, 203 genera, and 563 species. At the phylum level, microbiomes were dominated by Proteobacteria (mean relative abundance 71.6 ± 1.6%), Actinobacteria (16.6 ± 1.0%), and Firmicutes (6.7 ± 0.7% Figure 5C). Among the other phyla, we identified a smaller Archaea portion, primarily of the phylum Thaumarchaeota (2.4 ± 0.4%). At the family level, Proteobacteria was represented primarily by Bradyrhizobiaceae (6.6 ± 0.6%), Rhizobiaceae (33.9 ± 1.6%), and Sphingomonadaceae (8.8 ± 0.5%; Figure 5C). Within the phylum Actinobacteria, we identified Micrococcaceae (4.5 ± 0.5%), Mycobacteriaceae (3.3 ± 0.5%), and Streptomycetaceae (2.6 ± 0.2%). In Firmicutes, we identified only a single major family, Bacillaceae (6.6 ± 0.71%) that composed most of the abundance.

**Figure 5.**
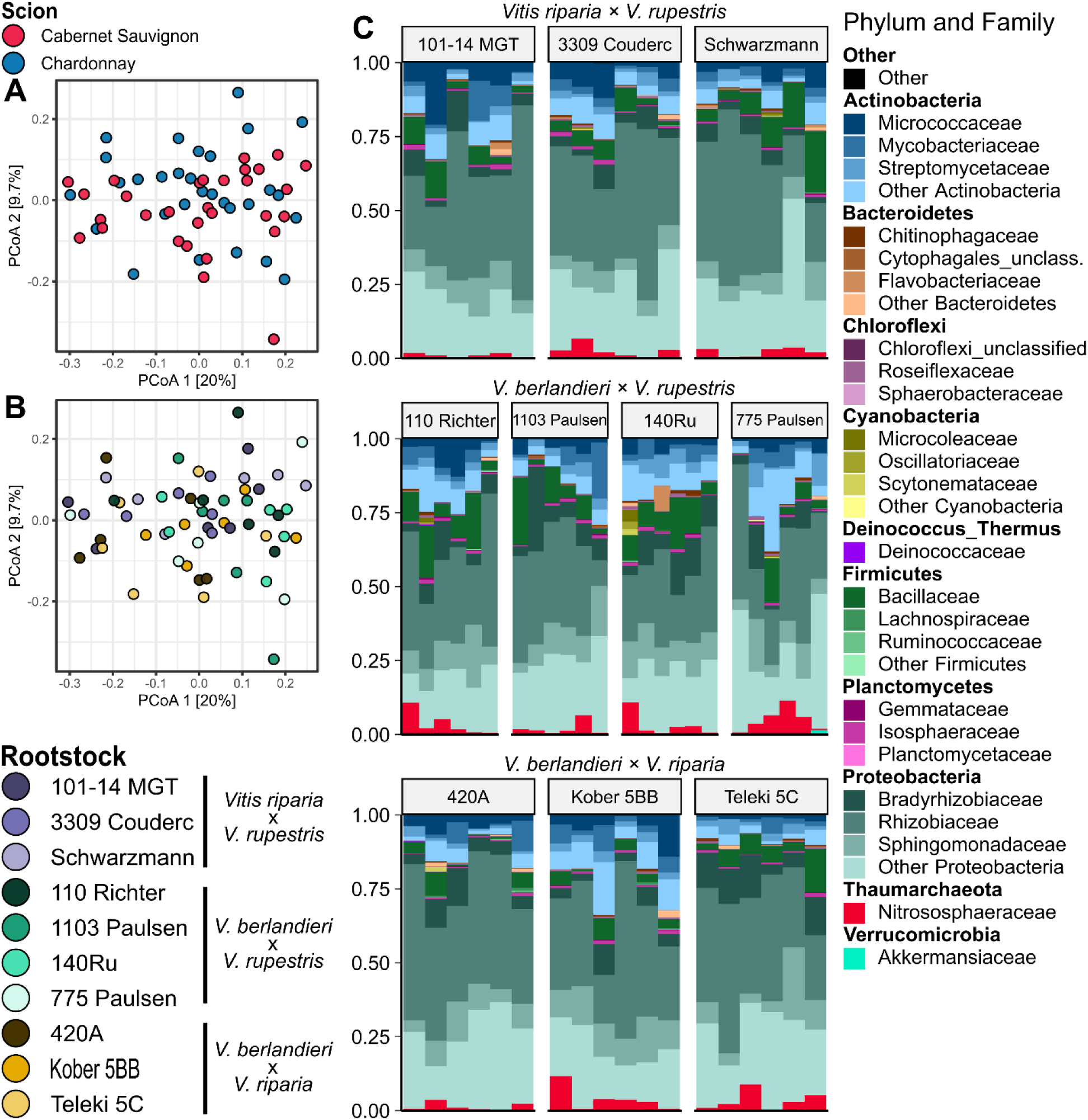
Rhizosphere microbiome taxonomic profiles. Principal coordinate analysis using Bray-Curtis Dissimilarity matrix with samples colored according to **A)** scion and **B)** rootstock genotypes. **C)** Taxonomic barplots showing the relative abundances of the top ten phyla, further subdivided into the top three families. Phyla outside the top ten are grouped into the “Other” category. Rootstock genotypes are separated by facets.

Few taxa were associated with a particular rootstock or scion genotype. In total, a single genus and four species exhibited differential abundance among genotypes, each with a unique rootstock/scion combination driving differentiation. Of these four species, *Pseudomonas moraviensis*, *Bacillus simplex*, and *Devosia insulae* were observed at low mean relative abundance (0.11%, 0.07%, and 0.03%, respectively) while *Bradyrhizobium ottawaense* was more abundant (1.92%; Figure S5A). For *Bradyrhizobium ottawaense*, ‘140Ru/Cabernet Sauvignon’ showed the highest mean relative abundance of any rootstock/scion combination (6.73%; Figure S5B). Similarly, *Pseudomonas moraviensis* was most abundant in ‘Teleki 5C/Chardonnay’ (0.63%; Figure S5A), *Bacillus simplex* in ‘110 Richter/Cabernet Sauvignon’ (0.51%), and *Devosia insulae* in ‘101-14 MGT/Cabernet Sauvignon’ (0.02%; Figure S5A). When grouping rootstocks by parentage, no taxa were associated with a particular group. Overall, taxonomic differences between rootstock/scion combinations are quite small, indicating that root systems recruit similar rhizosphere communities regardless of rootstock genotype and that scions do little to modulate the recruitment.

### Functional capacity of rhizosphere microbiomes is conserved across rootstock and scion genotypes

We assessed the functional capacity of the rhizosphere microbiome across rootstock and scion genotypes. We found the mean alignment rate of reads per sample to gene families was 14.5%. This was a result from multiple, non-mutually exclusive, factors including: the novelty and complexity of rhizosphere microbiome, lack of representation for many soil and rhizosphere associated microorganisms in protein databases, and conservative mapping settings. We identified 2,390,651 unique gene families (mean per sample = 334,747), which we fed into metabolic pathway predictions. We identified 493 unique metabolic pathways (mean per sample = 365) across the rhizosphere microbiome. These pathways were predominately in the super classes “Biosynthesis” (n = 276), “Degradation” (n = 201), and “Energy-Metabolism” (n = 65; note, some pathways are in multiple super classes). The “Degradation” super class was particularly enriched in the child classes “Aromatic-Compounds-Degradation” (n = 44), “Carbohydrates-Degradation” (n = 25), and “Amino-Acid-Degradation” (n = 21; note, some pathways share multiple child classes).

We identified metabolic pathways exhibiting differential enrichment across rootstock and scion genotypes. We found that only three metabolic pathways showed significant patterns of differential enrichment. These metabolic pathways were the superpathways “N-acetylglucosamine, N-acetylmannosamine and N-acetylneuraminate degradation” (BioCyc ID: GLCMANNANAUT-PWY), “pyrimidine deoxyribonucleotides de novo biosynthesis” (BioCyc ID: PWY-7211), and “flavin biosynthesis I” (BioCyc ID: RIBOSYN2-PWY). For both the GLCMANNANAUT-PWY and PWY-7211 pathways, the rootstock by scion interaction influenced enrichment (*p* < 0.01 and *p* = 0.01, respectively; Figure S6A). Post-hoc testing for these metabolic pathways showed that for GLCMANNANAUT-PWY no pairwise comparisons were significant after correction for multiple testing, whereas for PWY-7211 a single sample of the ‘Teleki 5C/Chardonnay’ was most responsible for the differential enrichment (Figure S6B). For the RIBOSYN2-PWY pathway, we observed a significant influence of rootstock genotype (ANOVA; “*rootstock” p* = 0.4, F_9,29_ = 2.30). Post-hoc testing revealed enrichment in the rootstock genotype ‘Teleki 5C’ in comparison to the base mean (*p* = 0.01; Figure 6). Modeling with parentage group, we found no pathways were differentially enriched.

**Figure 6.**
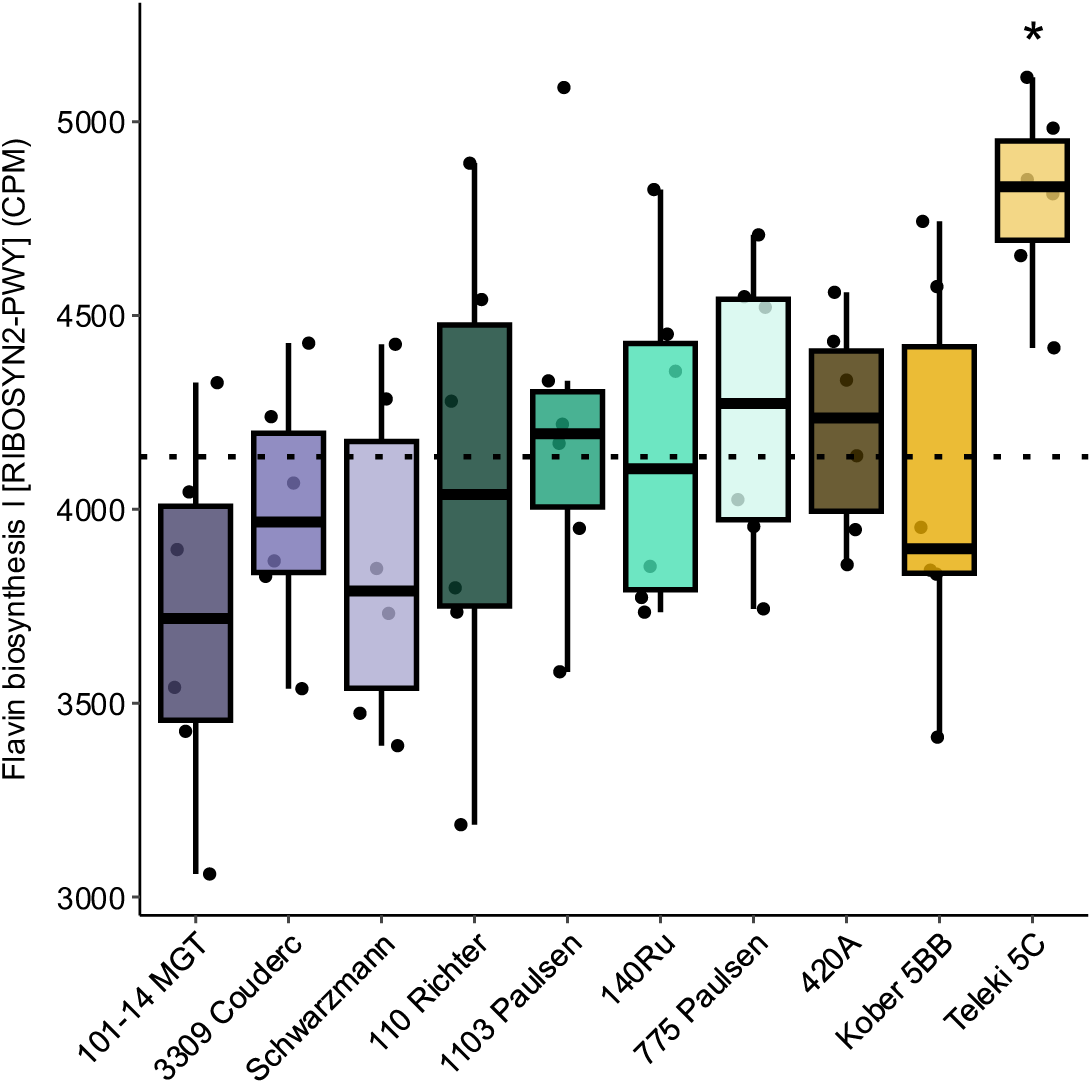
Flavin biosynthesis I exhibits differential enrichment for ‘Teleki 5C’. Flavin biosynthesis I (BioCyc ID: RIBOSYN2-PWY) pathway across rootstock genotypes with the base mean (dashed line). * denotes p < 0.01.

### Core taxa and functional capacity

In addition to identifying taxa and metabolic pathways were significantly influenced by rootstock and scion genotypes, we elected to define the core taxa and functional capacity of the grapevine rhizosphere. We selected taxa at the genus level and considered genera as core members if they met the following requirements: occupancy in 95% of samples (i.e., present in 57 of 60 samples) and a mean relative abundance of greater than 0.4% (80^th^ percentile; Figure S7). In total we identified 21 genera that met these conditions (Table S6). The identified core genera were distributed across five phyla; Proteobacteria (*n* = 11), Actinobacteria (*n* = 7), Firmicutes (*n* = 1), Chloroflexi (*n* = 1), and Planctomycetota (*n* = 1). We considered metabolic pathways as part of the core functional potential of grapevine rhizospheres if they met the following requirements, occupancy in 100% of samples and a mean enrichment of greater than 4215 CPM (80^th^ percentile; Figure S8). In total, 99 metabolic pathways met these conditions (Table S7). Of these 99 metabolic pathways, the largest MetaCyc hierarchy class was “Biosynthesis” which contained 83 of the pathways, followed by “Energy-Metabolism” with 13 pathways, and “Degradation” with 3 pathways. Breaking down the largest MetaCyc hierarchy, “Biosynthesis”, we found the majority of pathways were further separated into the subclasses “Amino-Acid-Biosynthesis” and “Nucleotide-Biosynthesis”.

### Elemental composition and its associations with the microbiome

As the root and rhizosphere microbiota are predominately enriched in taxa that are associated with nutrient uptake, we investigated the elemental composition of the rhizospheres. Due to the complex pairwise correlation structure of elemental compositions (Baxter, 2009) (Figure 2B), we elected to use principal component analysis (PCA) to decompose the data set into a smaller number of principal components (PC). The first ten PCs cumulatively capture 94.3% of the variance in elemental composition. For each of these ten PCs we fit a model with rootstock and scion genotype along with their interaction. We found that PC8 (2.22% of the variance) showed a significant influence of rootstock (*p* < 0.01) and PC6 (4.68% of the variance) showed a significant influence of scion (*p* < 0.01; Figure S9). Elements loading heavily on for PC8 were molybdenum (negative) and cadmium, potassium, iron (positive; Figure S9A). PC6 had strong negative loadings for cadmium and strong positive loadings for boron, selenium, iron and calcium (Figure S9B).

We then tested for associations between elemental composition PC6 and PC8 and bacterial genera and metabolic pathways from the rhizosphere microbiomes. Both PC6 and PC8 were significantly associated with a single genus (Figure 7A&B). PC8 was negatively correlated with the relative abundance of *Streptomyces* (Figure 7A), while PC6 was positively correlated with the relative abundance of *Mesorhizobium* (Figure 7B). For metabolic pathways we identified seven pathways with associations to PC6, however, P value correction rendered all associations non-significant (P_FDR_>0.05). Two pathways associated with arginine biosynthesis (BioCyc ID: ARGSYN-PWY and PWY-7400) had negative associations with PC6 (Figure 7C&D). Finally, we examined the remaining elemental composition PCs from the top ten that were not linked to scion or rootstock genotypes. and found no associations with any bacterial genera. In contrast, 51 metabolic pathways were associated with these additional elemental PCs (Figure S10).

**Figure 7.**
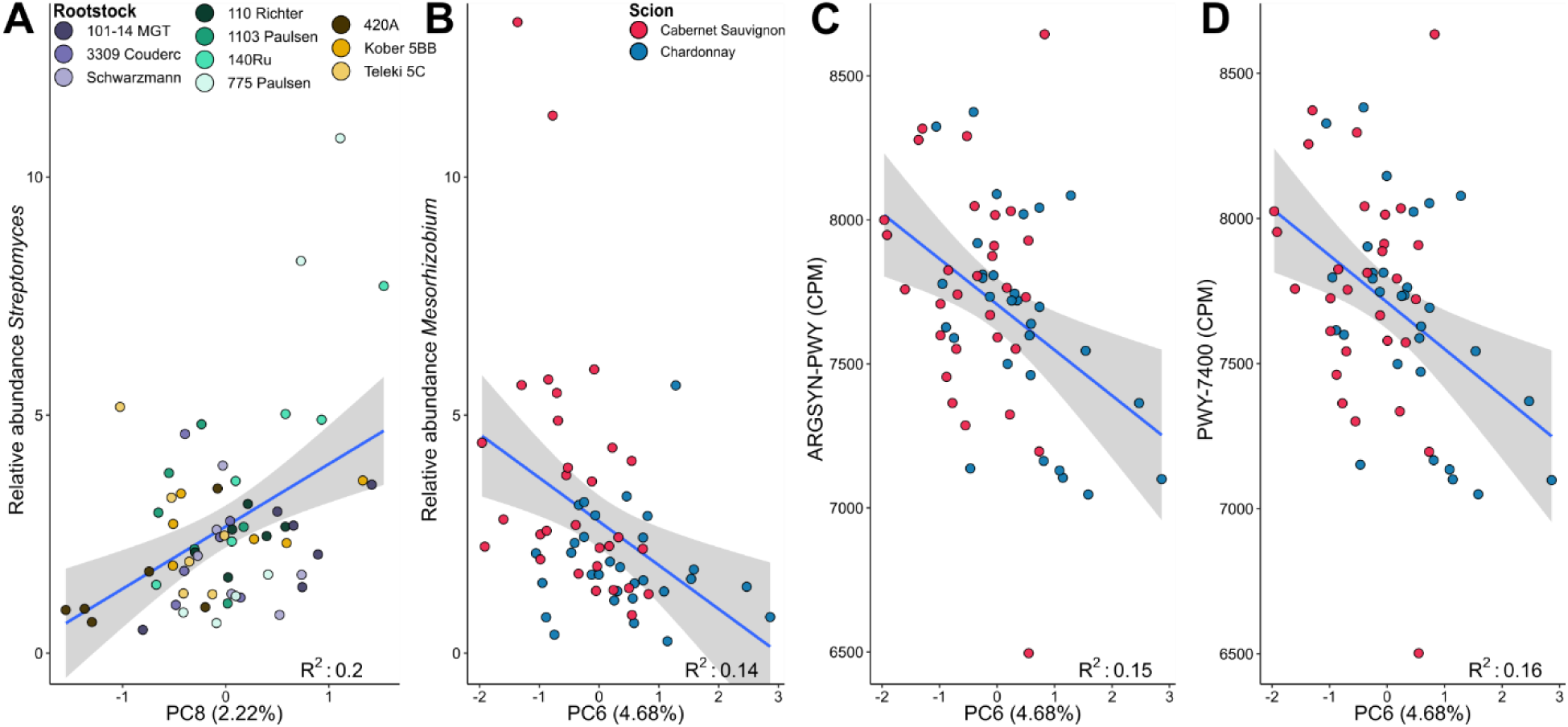
Associations between elemental composition principal components and the rhizosphere microbiome. PCA was used to decompose rhizosphere elemental profiles, and the resulting PCs were tested as response variables in models including rootstock, scion genotype, and their interaction. PCs with a significant effect of rootstock (PC8) or scion genotype (PC6; see Figure S9) were then examined for associations with the relative abundance of rhizosphere genera and metabolic pathways. **A)** *Streptomyces* was positively correlated with PC8 (linear regression; *p* < 0.01, F_1,58_ = 14.77, R^2^ = 0.2). **B)** *Mesorhizobium* was negatively correlated with PC6 (*p* = 0.03, F_1,58_ = 9.8, R^2^ = 0.14). Two arginine biosynthesis pathways were also associated with elemental composition PC6: **C)** L-arginine biosynthesis I (ARGSYN-PWY; *p* = 0.2, F_1,58_ = 10.38, R^2^ = 0.15) and **D)** L-arginine biosynthesis IV (PWY-7400; *p* = 0.2, F_1,58_ = 10.78, R^2^ = 0.16). Point fill indicates rootstock or scion genotype. Pathway enrichment values are given in copies per million (CPM), which account for gene length and the number of mapped reads.

## Discussion

Grapevine grafting is widespread in viticulture, yet researchers still lack a comprehensive understanding of how grafted rootstock genotypes and their associated microbiomes influence vine mineral nutrition. This study used a rootstock diversity trial in California’s Central Valley to survey a broad range of globally used rootstock genotypes, each grafted to two commercially important scions. We applied shotgun metagenomics and elemental composition analyses to identify links between the root microbiome and vine elemental composition.

Most research on grapevine mineral nutrition has focused on petioles or blades of fully developed leaves (Christensen, 1984; Benito *et al*., 2013; Schreiner & Scagel, 2017). These tissues are commonly utilized because they are easy to collect and reliably predict nutritional deficiencies and vine productivity. Although both tissues are informative, they have distinct physiological roles. Petioles function primarily as transport tissues within the shoot, whereas leaf blades are primarily responsible for photosynthesis and themselves exhibit complex source–sink dynamics (Yu *et al*., 2015). Rootstock genotypes have been shown to influence concentrations of various elements in both petioles and leaves (Harris *et al*., 2021, 2022; Migicovsky *et al*., 2021; Nikolaou *et al*., 2021), with rootstock genetic backgrounds (*i.e., V. berlandieri* and *V. rupestris*, and *V. riparia*) exhibiting characteristic patterns of elemental concentrations in these tissues (Wolpert *et al*., 2005; Gautier *et al*., 2020; Morel *et al*., 2024). Here, we tested patterns of elemental concentrations in roots relating individual rootstock genotypes, parentage groups, and genetic distance between rootstocks. We found that specific genotypes showed significant associations with individual elements, but these associations were idiosyncratic to parentage groups, and genetic distance among rootstock genotypes was not related to similarity in elemental composition in either tissue. This pattern may reflect the tissues selected for this study, roots and berries. However, because roots and berries represent source and sink tissues, respectively, our results suggest that rootstock genotypes have only a minimal influence on whole-vine elemental composition at this vineyard. Surveys of the elemental composition of these tissues across a larger number of vineyards or regions, similar to those conducted for petioles (Wolpert *et al*., 2005), would help assess the environmental contribution to this G×E interaction.

The rhizosphere microbiomes of different rootstock/scion combinations were similar. This is consistent with previous amplicon-based sequencing studies that illustrated a small role for rootstock genotype in shaping rhizosphere microbiomes (D’Amico *et al*., 2018; Marasco *et al*., 2018; Berlanas *et al*., 2019; Swift *et al*., 2021, 2023; Wright *et al*., 2022, p. 20). As root exudates serve as a primary mechanism for selection of the plant rhizosphere (Haichar *et al*., 2008; Sasse *et al*., 2018), our results suggest that root exudation profiles are similar across rootstock genotypes. Alternatively, since our collections focused on a single vineyard, variation in soil topography or other environmental conditions in this natural setting could outweigh any subtle differences in root exudate profiles. Indeed, prior work has shown that soil type can override host-selection effects across a broad phylogenetic breadth of plants (Yeoh *et al*., 2017). Nevertheless, since our study focused on a single location and because previous 16S rRNA gene sequencing of soils and roots from this vineyard revealed little intra-site heterogeneity in bacterial communities (Swift *et al*., 2023), it is plausible that root exudation profiles are relatively consistent across these rootstock genotypes

The microbial taxa we identified as variable across rootstock/scion combinations are notable in their patterns and putative plant interactions. At the species level, *Bradyrhizobium ottawaense* and *Devosia insulae* were among the most differentially abundant taxa.

*Bradyrhizobium ottawaense* was influenced by the interaction of rootstock and scion genotypes (Figure S5A). This bacterium is capable of nitrogen fixation and was originally isolated from soybean nodules (Yu *et al*., 2014; Nguyen *et al*., 2018). As nodule formation is generally restricted to leguminous plant species (van Rhijn & Vanderleyden, 1995; Sprent, 2007), the taxon identified as *B. ottawaense* in our dataset is likely a closely related species that is not currently represented in proteomic databases. Nevertheless, members of this genus have also been recovered in non-leguminous hosts, including sorghum and *Arabidopsis thaliana* (Schneijderberg *et al*., 2018; Wasai-Hara *et al*., 2020). *Devosia insulae* only made up a small proportion of the rhizosphere microbiome but was enriched in ‘101-14 MGT/Cabernet Sauvignon’ (Figure S5B). *Devosia* spp. have typically been isolated from soils but occupy a wide number of ecological niches (Talwar *et al*., 2020). For example, five *Devosia* isolates were recovered from *Arabidopsis thaliana* roots grown in natural soils (Bai *et al*., 2015). A phylogenetic study of *Devosia* found *D. insulae*, originally isolated from soil (Yoon *et al*., 2007), formed a monophyletic clade with several other isolates of agricultural soil or plant rhizosphere origin (Talwar *et al*., 2020). The ubiquity of this genus in plant rhizospheres (Mohd Nor *et al*., 2017; Wolters *et al*., 2018; Walters *et al*., 2018; Wang *et al*., 2022) and the ability of some members of the genus to produce phytohormones (Chhetri *et al*., 2022) suggests plant-growth promoting properties of this genus that warrants further study.

Similar to the taxonomic profiles, few metabolic pathways differed between rootstock/scion combinations, indicating a common functional repertoire for grafted grapevines at this location, consistent with results from other plant systems (Jin *et al*., 2017; Xu *et al*., 2018b). Of the three predicted pathways that were differentially enriched, we found that “flavin biosynthesis I” (BioCyc ID: RIBOSYN2-PWY) was significantly enriched in the rootstock genotype ‘Teleki 5C’ independent of scion (Figure 6). A key product of this metabolic pathway is riboflavin (also known as vitamin B2), a molecule with numerous roles in plant growth stimulation (Dakora *et al*., 2015). Riboflavin is released by rhizosphere microorganisms (Yurgel *et al*., 2014) and can be rapidly degraded by other microorganisms to lumichrome (*Pseudomonas* spp. (Yanagita & Foster, 1956; Phillips *et al*., 1999). Lumichrome and riboflavin supplementation in plants have shown increases in root respiration and growth (Matiru & Dakora, 2005), disease resistance (Dong & Beer, 2000), photosynthetic rate (Khan *et al*., 2008), and starch production (Gouws *et al*., 2012). These findings highlight the need to explore how rootstock genotype may influence this expression of this pathway.

The diversity of rootstocks in this study provides a unique opportunity to identify the core *Vitis* rhizosphere microbiome and its associated metabolic pathways. Core features of a microbiome are most often defined using two metrics: occupancy (frequency of occurrence) and relative abundance (Shade & Stopnisek, 2019; Risely, 2020; Neu *et al*., 2021). We chose conservative filtering settings to reduce noise. The most abundant genera conserved were predominately diazotrophic, possibly due to their role in supporting plant nutrition via nitrogen fixation. *Rhizobium* was the top genus defined as core in terms of its position on the abundance–occurrence curve (Table S6), with a mean relative abundance of ∼30% and occupancy in 100% of samples. *Rhizobium* is associated with many plant growth promoting properties (Antoun *et al*., 1998; Mehboob *et al*., 2009) and is consistently identified as a member of core rhizosphere microbiomes across plant species (Lundberg *et al*., 2012; Bulgarelli *et al*., 2012; Jin *et al*., 2017; Walters *et al*., 2018; Xu *et al*., 2018b; Chen *et al*., 2019). Many of the core genera we identified have been linked to plant growth promotion, including *Sphingomonas*, which stimulates lateral root and root hair formation in *Arabidopsis* (Luo *et al*., 2019); *Bacillus*, which induces systemic resistance to phytopathogens across plant species (Kloepper *et al*., 2004; Cazorla *et al*., 2007; Mercado-Flores *et al*., 2014); and *Variovorax*, which enhances tolerance to abiotic stress (Jiang *et al*., 2012). Together, these genera likely contribute to the plant growth–promoting functions of the grapevine rhizosphere.

We utilized similar filtering in order to identify core metabolic functions of the grapevine rhizosphere. Core functional capacities of microbial communities are predicted to be more tightly conserved than taxonomic profiles (Louca *et al*., 2016, 2018; Lemanceau *et al*., 2017). We found support for the conservation of a functional core containing many metabolic pathways present across all the samples when filtered at the 80th percentile of count depth (CPM; Table S6). The core metabolic pathways were predominately associated with the MetaCyc subclasses “Amino-Acid-Biosynthesis” and “Nucleotide-Biosynthesis”. These subclasses include pathways essential survival by the synthesis of RNA, DNA, and amino acids. A metagenomic survey of the perennial fruit crop genus *Citrus* found that compared to bulk soil the rhizosphere samples were depleted in KEGG Orthologs related to amino acid and nucleotide synthesis (Xu *et al*., 2018b). As plants release carbon-rich exudates that include amino acids (Jones *et al*., 2009; Sasse *et al*., 2018), this may reduce microbial investment in amino acid and nucleotide biosynthesis (Zhalnina *et al*., 2018). However, amino acid synthesis has been shown to be requirement for colonization of plant roots for some microorganisms (Simons *et al*., 1997) and this functionality is recovered in other rhizosphere metagenomic studies (Jin *et al*., 2017; Chen *et al*., 2022). Our analysis revealed that these functional capacities are highly conserved in the rhizosphere community, but we cannot assess whether they are depleted relative to bulk soil in this vineyard.

We found that *Streptomyces* and *Mesorhizobium* were significantly associated with root elemental profiles. Additionally, two arginine biosynthesis gene pathways were negatively correlated with these profiles, though they were not significant after multiple testing correction (Figure 7). *Streptomyces* was positively correlated with elemental PC8, for which rootstock explained a significant amount of variance (Figure 7A). This genus is known to include many species with plant growth-promoting properties (Viaene *et al*., 2016; Rey & Dumas, 2017) and can solubilize some nutrients *in vitro* (Yagi *et al*., 1971; Gadkari *et al*., 1992; Gupta *et al*., 2010; Jog *et al*., 2014). However, validation of this functionality *in planta* is lacking but see (Rungin *et al*., 2012). *Mesorhizobium* was negatively correlated with a PC that was significantly explained by scion genotype (Figure 7B). This genus is typically associated with the nodules of leguminous plant species (Nguyen *et al*., 2015; Meng *et al*., 2022; Knežević *et al*., 2022)but is also frequently detected in rhizospheres of non-legumes (Chaparro *et al*., 2014; Qiao *et al*., 2017; Yeoh *et al*., 2017; Zhalnina *et al*., 2018; Ling *et al*., 2022), including grapevines (Marasco *et al*., 2013; Andreolli *et al*., 2017). *Mesorhizobium* spp. have been shown to produce phytohormones as well as solubilize potassium and phosphate, and improved plant growth when inoculated in tomato (Menéndez *et al*., 2020). In that study, shoots of the inoculated *vs.* control tomato plants differed in concentrations of multiple elements including iron and calcium which loaded negatively on PC6 in this study (Figure 7B-D). Both genera warrant further investigation to clarify their responses to rootstock and scion genotype, as well as their broader roles in shaping plant elemental profiles.

## Conclusion

The elemental composition of grapevine tissues is a critical metric for viticulturists when guiding management decisions, underscoring the importance of rootstock selection. We found that only a small number of root element concentrations were influenced by rootstock genotype, and this effect was minor and unrelated to genetic relatedness among rootstocks. For berry element concentrations we found patterns that were dictated primarily by the scion cultivar. The rhizosphere microbiome composition was similarly conserved across the diversity of rootstocks, indicating that rootstocks recruit a consistent set of microorganisms performing similar functions in this environment. This study lays the groundwork for defining the core *Vitis* rhizosphere microbiome, both taxonomically and functionally, through surveys of diverse commercial rootstock genotypes that represent the majority of vineyards worldwide.

## Data availability

Analysis code is available in the GitHub repository associated with this study https://github.com/Kenizzer/grafted_grapevine_microbiome. All raw sequencing data are available in the NCBI Short Read Archive under BioProject ID PRJNA1367536.

## Acknowledgments

We thank Peter Cousins (Gallo Winery) for logistical support through all phases of collections and vineyard managers and staff Danielle Hopkins, Jorge Fuentes, Keith Striegler, Lindsay Jordan, and Octavio Viveros for their support. In addition, we thank Daniel H. Chitwood (Michigan State University) and Maggie R. Wagner (University of Kansas) for helpful discussions in preparing this article; Niyati Bhakta and Leah Pinkner for assistance with sample preparation. This project was made possible in part by funds supporting a summer training program in which undergraduate students from partnering institutions contributed to field work in commercial vineyards in California. We acknowledge the following students for their help in this work: Julie Curless (Missouri State University) and Zach Helget (South Dakota State University). This work was funded by the National Science Foundation Plant Genome Research Program under Award No. 1546869. JFS was supported by an NSF Graduate Research Fellowship under Grant No. 1758713 and NSF Postdoctoral Research Fellowship under Award No. 2305703. ZM was supported by funding from a Natural Sciences and Engineering Research Council (NSERC) of Canada Discovery Grant and the Canada Research Chairs Program (CRC-2021-00314).

## Author Contributions

JFS, ZM, and AJM designed the experimental and collection design. JFS and ZM performed sample collection. JFS performed sample processing. JFS, ZM, and ZNH performed data analysis. All authors contributed to data interpretation, the writing and reviewing of the manuscript, and approved the final draft.

## References

Adamack AT, Gruber B. 2014. PopGenReport: simplifying basic population genetic analyses in R. Methods in Ecology and Evolution 5: 384–387.

Andreolli M, Lampis S, Vallini G. 2017. Diversity, Distribution and Functional Role of Bacterial Endophytes in Vitis vinifera. In: Endophytes: Biology and Biotechnology. 1–35.

Antoun H, Beauchamp CJ, Goussard N, Chabot R, Lalande R. 1998. Potential of Rhizobium and Bradyrhizobium species as plant growth promoting rhizobacteria on non-legumes: Effect on radishes (Raphanus sativus L.). Plant and Soil 204: 57–67.

Bai Y, Müller DB, Srinivas G, Garrido-Oter R, Potthoff E, Rott M, Dombrowski N, Münch PC, Spaepen S, Remus-Emsermann M, et al. 2015. Functional overlap of the Arabidopsis leaf and root microbiota. Nature 528: 364–369.

Bates D, Mächler M, Bolker B, Walker S. 2015. Fitting Linear Mixed-Effects Models Using lme4. Journal of Statistical Software 67.

Bavaresco L, Giachino E, Pezzutto S. 2003. Grapevine rootstock effects on lime-induced chlorosis, nutrient uptake, and source-sink relationships. Journal of Plant Nutrition 26: 1451–1465.

Baxter I. 2009. Ionomics: studying the social network of mineral nutrients. Current Opinion in Plant Biology 12: 381–386.

Baxter I. 2010. Ionomics: The functional genomics of elements. Briefings in Functional Genomics and Proteomics 9: 149–156.

Beghini F, McIver LJ, Blanco-Míguez A, Dubois L, Asnicar F, Maharjan S, Mailyan A, Manghi P, Scholz M, Thomas AM, et al. 2021. Integrating taxonomic, functional, and strain-level profiling of diverse microbial communities with bioBakery 3. eLife 10: 1–42.

Benito A, Romero I, Domínguez N, García-Escudero E, Martín I. 2013. Leaf blade and petiole analysis for nutrient diagnosis in Vitis vinifera L. cv. Garnacha tinta. Australian Journal of Grape and Wine Research 19: 285–298.

Berlanas C, Berbegal M, Elena G, Laidani M, Cibriain JF, Sagües A, Gramaje D. 2019. The Fungal and Bacterial Rhizosphere Microbiome Associated With Grapevine Rootstock Genotypes in Mature and Young Vineyards. Frontiers in Microbiology 10: 1–16.

Bolger AM, Lohse M, Usadel B. 2014. Trimmomatic: a flexible trimmer for Illumina sequence data. Bioinformatics 30: 2114–2120.

Bonkowski M. 2004. Protozoa and plant growth: the microbial loop in soil revisited. New Phytologist 162: 617–631.

Bulgarelli D, Rott M, Schlaeppi K, Van Themaat EVL, Ahmadinejad N, Assenza F, Rauf P, Huettel B, Reinhardt R, Schmelzer E, et al. 2012. Revealing structure and assembly cues for Arabidopsis root-inhabiting bacterial microbiota. Nature 488: 91–5.

Canaguier A, Grimplet J, Di Gaspero G, Scalabrin S, Duchêne E, Choisne N, Mohellibi N, Guichard C, Rombauts S, Le Clainche I, et al. 2017. A new version of the grapevine reference genome assembly (12X.v2) and of its annotation (VCost.v3). Genomics Data 14: 56–62.

Cazorla FM, Romero D, Pérez-García A, Lugtenberg BJJ, Vicente AD, Bloemberg G. 2007. Isolation and characterization of antagonistic Bacillus subtilis strains from the avocado rhizoplane displaying biocontrol activity. Journal of Applied Microbiology 103: 1950–1959.

Chaparro JM, Badri D V., Vivanco JM. 2014. Rhizosphere microbiome assemblage is affected by plant development. ISME Journal 8: 790–803.

Chen C, Wang M, Zhu J, Tang Y, Zhang H, Zhao Q, Jing M, Chen Y, Xu X, Jiang J, et al. 2022. Long-term effect of epigenetic modification in plant–microbe interactions: modification of DNA methylation induced by plant growth-promoting bacteria mediates promotion process. Microbiome 10: 1–19.

Chen L, Xin X, Zhang J, Redmile-Gordon M, Nie G, Wang Q. 2019. Soil Characteristics Overwhelm Cultivar Effects on the Structure and Assembly of Root-Associated Microbiomes of Modern Maize. Pedosphere 29: 360–373.

Chhetri G, Kim I, Kang M, Kim J, So Y, Seo T. 2022. Devosia rhizoryzae sp. nov., and Devosia oryziradicis sp. nov., novel plant growth promoting members of the genus Devosia, isolated from the rhizosphere of rice plants. Journal of Microbiology 60: 1–10.

Christensen P. 1984. Nutrient Level Comparisons of Leaf Petioles and Blades in Twenty-Six Grape Cultivars Over Three Years (1979 through 1981). American Journal of Enology and Viticulture 35: 124–133.

Dai Z, Liu G, Chen H, Chen C, Wang J, Ai S, Wei D, Li D, Ma B, Tang C, et al. 2020. Long-term nutrient inputs shift soil microbial functional profiles of phosphorus cycling in diverse agroecosystems. The ISME Journal 14: 757–770.

Dakora FD, Matiru VN, Kanu AS. 2015. Rhizosphere ecology of lumichrome and riboflavin, two bacterial signal molecules eliciting developmental changes in plants. Frontiers in Plant Science 6: 1–11.

D’Amico F, Candela M, Turroni S, Biagi E, Brigidi P, Bega A, Vancini D, Rampelli S. 2018. The rootstock regulates microbiome diversity in root and rhizosphere compartments of Vitis vinifera cultivar Lambrusco. Frontiers in Microbiology 9: 2240.

Dong H, Beer SV. 2000. Riboflavin induces disease resistance in plants by activating a novel signal transduction pathway. Phytopathology 90: 801–811.

Ewels P, Magnusson M, Lundin S, Käller M. 2016. MultiQC: summarize analysis results for multiple tools and samples in a single report. Bioinformatics 32: 3047–3048.

Gadkari D, Mörsdorf G, Meyer O. 1992. Chemolithoautotrophic assimilation of dinitrogen by Streptomyces thermoautotrophicus UBT1: identification of an unusual N2-fixing system. Journal of Bacteriology 174: 6840–6843.

Gautier A, Cookson SJ, Lagalle L, Ollat N, Marguerit E. 2020. Influence of the three main genetic backgrounds of grapevine rootstocks on petiolar nutrient concentrations of the scion, with a focus on phosphorus. OENO One 54: 1–13.

Gouws LM, Botes E, Wiese AJ, Trenkamp S, Torres-Jerez I, Tang Y, Hills PN, Usadel B, Lloyd JR, Fernie AR, et al. 2012. The Plant Growth Promoting Substance, Lumichrome, Mimics Starch, and Ethylene-Associated Symbiotic Responses in Lotus and Tomato Roots. Frontiers in Plant Science 3: 1–20.

Gupta N, Sahoo D, Chand Basak U. 2010. Evaluation of in vitro solubilization potential of phosphate solubilising Streptomyces isolated from phyllosphere of Heritiera fomes (mangrove). African Journal of Microbiology Research 4: 136–142.

Haichar FEZ, Marol C, Berge O, Rangel-Castro JI, Prosser JI, Balesdent J, Heulin T, Achouak W. 2008. Plant host habitat and root exudates shape soil bacterial community structure. The ISME Journal 2: 1221–1230.

Harris ZN, Awale M, Bhakta N, Chitwood DH, Fennell A, Frawley E, Klein LL, Kovacs LG, Kwasniewski M, Londo JP, et al. 2021. Multi-dimensional leaf phenotypes reflect root system genotype in grafted grapevine over the growing season. GigaScience 10: 1–17.

Harris ZN, Pratt JE, Bhakta N, Frawley E, Klein LL, Kwasniewski MT, Migicovsky Z, Miller AJ. 2022. Temporal and environmental factors interact with rootstock genotype to shape leaf elemental composition in grafted grapevines. Plant Direct 6: e440.

Harris ZN, Pratt JE, Kovacs LG, Klein LL, Kwasniewski MT, Londo JP, Wu AS, Miller AJ. 2023. Grapevine scion gene expression is driven by rootstock and environment interaction. BMC Plant Biology 23: 211.

Harrison XA, Donaldson L, Correa-Cano ME, Evans J, Fisher DN, Goodwin CED, Robinson BS, Hodgson DJ, Inger R. 2018. A brief introduction to mixed effects modelling and multi-model inference in ecology. PeerJ 6: e4794.

Ibacache AG, Sierra CB. 2009. Influence of Rootstocks on Nitrogen, Phosphorus and Potassium Content in Petioles of Four Table Grape Varieties. Chilean journal of agricultural research 69: 503–508.

Jacoby R, Peukert M, Succurro A, Koprivova A, Kopriva S. 2017. The Role of Soil Microorganisms in Plant Mineral Nutrition—Current Knowledge and Future Directions. Frontiers in Plant Science 8: 1–19.

Jia B, Chang X, Fu Y, Heng W, Ye Z, Liu P, Liu L, Al Shoffe Y, Watkins CB, Zhu L. 2022. Metagenomic analysis of rhizosphere microbiome provides insights into occurrence of iron deficiency chlorosis in field of Asian pears. BMC Microbiology 22: 1–16.

Jiang F, Chen L, Belimov AA, Shaposhnikov AI, Gong F, Meng X, Hartung W, Jeschke DW, Davies WJ, Dodd IC. 2012. Multiple impacts of the plant growth-promoting rhizobacterium Variovorax paradoxus 5C-2 on nutrient and ABA relations of Pisum sativum. Journal of Experimental Botany 63: 6421–6430.

Jin T, Wang Y, Huang Y, Xu J, Zhang P, Wang N, Liu X, Chu H, Liu G, Jiang H, et al. 2017. Taxonomic structure and functional association of foxtail millet root microbiome. GigaScience 6: 1–12.

Jog R, Pandya M, Nareshkumar G, Rajkumar S. 2014. Mechanism of phosphate solubilization and antifungal activity of Streptomyces spp. isolated from wheat roots and rhizosphere and their application in improving plant growth. Microbiology 160: 778–788.

Jones DL, Nguyen C, Finlay RD. 2009. Carbon flow in the rhizosphere: Carbon trading at the soil-root interface. Plant and Soil 321: 5–33.

Kassambara A. 2023. ggpubr.

Kassambara A, Mundt F. 2020. factoextra: Extract and Visualize the Results of Multivariate Data Analyses.

Khan W, Prithiviraj B, Smith DL. 2008. Nod factor [Nod Bj V (C18:1, MeFuc)] and lumichrome enhance photosynthesis and growth of corn and soybean. Journal of Plant Physiology 165: 1342–1351.

Kloepper JW, Ryu C-M, Zhang S. 2004. Induced Systemic Resistance and Promotion of Plant Growth by Bacillus spp. Phytopathology® 94: 1259–1266.

Knežević M, Berić T, Buntić A, Jovković M, Avdović M, Stanković S, Delić D, Stajković-Srbinović O. 2022. Native Mesorhizobium strains improve yield and nutrient composition of the common bird’s-foot trefoil grown in an acid soil. Rhizosphere 21: 100487.

Kosman E, Leonard KJ. 2005. Similarity coefficients for molecular markers in studies of genetic relationships between individuals for haploid, diploid, and polyploid species. Molecular Ecology 14: 415–424.

Krithika S, Balachandar D. 2016. Expression of zinc transporter genes in rice as influenced by zinc-solubilizing enterobacter cloacae strain ZSB14. Frontiers in Plant Science 7: 1–9.

Langmead B, Salzberg SL. 2012. Fast gapped-read alignment with Bowtie 2. Nature methods 9: 357–9.

Lecourt J, Lauvergeat V, Ollat N, Vivin P, Cookson SJ. 2015. Shoot and root ionome responses to nitrate supply in grafted grapevines are rootstock genotype dependent. Australian Journal of Grape and Wine Research 21: 311–318.

Lemanceau P, Blouin M, Muller D, Moënne-Loccoz Y. 2017. Let the Core Microbiota Be Functional. Trends in Plant Science 22: 583–595.

Lenth R, Love J, Herve M. 2020. emmeans: Estimated Marginal Means, aka Least-Squares Means. : https://cran.r-project.org/package=emmeans.

Ling N, Wang T, Kuzyakov Y. 2022. Rhizosphere bacteriome structure and functions. Nature Communications 13: 1–13.

Louca S, Jacques SMS, Pires APF, Leal JS, Srivastava DS, Parfrey LW, Farjalla VF, Doebeli M. 2016. High taxonomic variability despite stable functional structure across microbial communities. Nature Ecology & Evolution 1: 1–12.

Louca S, Polz MF, Mazel F, Albright MBN, Huber JA, O’Connor MI, Ackermann M, Hahn AS, Srivastava DS, Crowe SA, et al. 2018. Function and functional redundancy in microbial systems. Nature Ecology & Evolution 2: 936–943.

Lundberg DS, Lebeis SL, Paredes SH, Yourstone S, Gehring J, Malfatti S, Tremblay J, Engelbrektson A, Kunin V, del Rio TG, et al. 2012. Defining the core Arabidopsis thaliana root microbiome. Nature 488: 86–90.

Luo Y, Wang F, Huang Y, Zhou M, Gao J, Yan T, Sheng H, An L. 2019. Sphingomonas sp. Cra20 increases plant growth rate and alters rhizosphere microbial community structure of Arabidopsis thaliana under drought stress. Frontiers in Microbiology 10.

Mallick H, Rahnavard A, McIver LJ, Ma S, Zhang Y, Nguyen LH, Tickle TL, Weingart G, Ren B, Schwager EH, et al. 2021. Multivariable association discovery in population-scale meta-omics studies. PLOS Computational Biology 17: e1009442.

Marasco R, Alturkey H, Fusi M, Brandi M, Ghiglieno I, Valenti L, Daffonchio D. 2022. Rootstock–scion combination contributes to shape diversity and composition of microbial communities associated with grapevine root system. Environmental Microbiology 00.

Marasco R, Rolli E, Fusi M, Cherif A, Abou-Hadid A, El-Bahairy U, Borin S, Sorlini C, Daffonchio D. 2013. Plant Growth Promotion Potential Is Equally Represented in Diverse Grapevine Root-Associated Bacterial Communities from Different Biopedoclimatic Environments. BioMed Research International 2013: 1–17.

Marasco R, Rolli E, Fusi M, Michoud G, Daffonchio D. 2018. Grapevine rootstocks shape underground bacterial microbiome and networking but not potential functionality. Microbiome 6: 3.

Matiru VN, Dakora FD. 2005. Xylem transport and shoot accumulation of lumichrome, a newly recognized rhizobial signal, alters root respiration, stomatal conductance, leaf transpiration and photosynthetic rates in legumes and cereals. New Phytologist 165: 847–855.

Mclver LJ, Abu-Ali G, Franzosa EA, Schwager R, Morgan XC, Waldron L, Segata N, Huttenhower C. 2018. bioBakery: a meta’omic analysis environment. Bioinformatics 34: 1235–1237.

McMurdie PJ, Holmes S. 2013. phyloseq: An R Package for Reproducible Interactive Analysis and Graphics of Microbiome Census Data. PLoS ONE 8: e61217.

Mehboob I, Naveed M, Zahir ZA. 2009. Rhizobial Association with Non-Legumes: Mechanisms and Applications. Critical Reviews in Plant Sciences 28: 432–456.

Menéndez E, Pérez-Yépez J, Hernández M, Rodríguez-Pérez A, Velázquez E, León-Barrios M. 2020. Plant Growth Promotion Abilities of Phylogenetically Diverse Mesorhizobium Strains: Effect in the Root Colonization and Development of Tomato Seedlings. Microorganisms 8: 412.

Meng D, Liu Y-L, Zhang J-J, Gu P-F, Fan X-Y, Huang Z-S, Ji Y, Meng H, Du Z-J, Li W-M, et al. 2022. Mesorhizobium xinjiangense sp. nov., isolated from rhizosphere soil of Alhagi sparsifolia. Archives of Microbiology 204: 29.

Mercado-Flores Y, Cárdenas-Álvarez IO, Rojas-Olvera AV, Pérez-Camarillo JP, Leyva-Mir SG, Anducho-Reyes MA. 2014. Application of Bacillus subtilis in the biological control of the phytopathogenic fungus Sporisorium reilianum. Biological Control 76: 36–40.

Migicovsky Z, Cousins P, Jordan LM, Myles S, Striegler RK, Verdegaal P, Chitwood DH. 2021. Grapevine rootstocks affect growth-related scion phenotypes. Plant Direct 5: e00324.

Migicovsky Z, Harris ZN, Klein LL, Li M, McDermaid A, Chitwood DH, Fennell A, Kovacs LG, Kwasniewski M, Londo JP, et al. 2019. Rootstock effects on scion phenotypes in a ‘Chambourcin’ experimental vineyard. Horticulture Research 6: 64.

Mohd Nor MN, Sabaratnam V, Tan GYA. 2017. Devosia elaeis sp. nov., isolated from oil palm rhizospheric soil. International Journal of Systematic and Evolutionary Microbiology 67: 851–855.

Morel M, Cookson SJ, Costa J-PD, Ollat N, Marguerit E. 2024. The role of rootstock and its genetic background in plant mineral status: the relationship between petiole analyses and deficiency symptoms. OENO One 58.

Mudge K, Janick J, Scofield S, Goldschmidt EE. 2009. A History of Grafting. In: Horticultural Reviews. Hoboken, NJ, USA: John Wiley & Sons, Inc., 437–493.

Naylor D, Coleman-Derr D. 2018. Drought Stress and Root-Associated Bacterial Communities. Frontiers in Plant Science 8: 1–16.

Naylor D, DeGraaf S, Purdom E, Coleman-Derr D. 2017. Drought and host selection influence bacterial community dynamics in the grass root microbiome. The ISME Journal 11: 2691–2704.

Neu AT, Allen EE, Roy K. 2021. Defining and quantifying the core microbiome: Challenges and prospects. Proceedings of the National Academy of Sciences 118: 1–10.

Nguyen HDT, Cloutier S, Bromfield ESP. 2018. Complete Genome Sequence of Bradyrhizobium ottawaense OO99 T, an Efficient Nitrogen-Fixing Symbiont of Soybean (FJ Stewart, Ed.). Microbiology Resource Announcements 7: 1–2.

Nguyen TM, Pham VHT, Kim J. 2015. Mesorhizobium soli sp. nov., a novel species isolated from the rhizosphere of Robinia pseudoacacia L. in South Korea by using a modified culture method. Antonie van Leeuwenhoek 108: 301–310.

Nikolaou K-E, Chatzistathis T, Theocharis S, Argiriou A, Koundouras S, Zioziou E, Nikolaou K-E, Chatzistathis T, Theocharis S, Argiriou A, et al. 2021. Effects of Salinity and Rootstock on Nutrient Element Concentrations and Physiology in Own-Rooted or Grafted to 1103 P and 101-14 Mgt Rootstocks of Merlot and Cabernet Franc Grapevine Cultivars under Climate Change. Sustainability 13.

Oksanen J, Guillaume Blanchet F, Friendly M, Kindt R, Legendre P, McGlinn D, Minchin PR, O’Hara RB, Simpson GL, Solymos P, et al. 2019. vegan: Community Ecology Package.

Ollat N, Bordenave L, Tandonnet JP, Boursiquot JM, Marguerit E. 2016. Grapevine rootstocks: origins and perspectives. Acta Horticulturae: 11–22.

Phillips DA, Joseph CM, Yang G-P, Martínez-Romero E, Sanborn JR, Volpin H. 1999. Identification of lumichrome as a Sinorhizobium enhancer of alfalfa root respiration and shoot growth. Proceedings of the National Academy of Sciences 96: 12275–12280.

Qiao Q, Wang F, Zhang J, Chen Y, Zhang C, Liu G, Zhang H, Ma C, Zhang J. 2017. The Variation in the Rhizosphere Microbiome of Cotton with Soil Type, Genotype and Developmental Stage. Scientific Reports 7: 1–10.

R Core Team. 2025. R: A Language and Environment for Statistical Computing.

Rey T, Dumas B. 2017. Plenty Is No Plague: Streptomyces Symbiosis with Crops. Trends in Plant Science 22: 30–37.

van Rhijn P, Vanderleyden J. 1995. The Rhizobium-plant symbiosis. Microbiological reviews 59: 124–142.

Riaz S, Pap D, Uretsky J, Laucou V, Boursiquot J-M, Kocsis L, Andrew Walker M. 2019. Genetic diversity and parentage analysis of grape rootstocks. Theoretical and Applied Genetics.

Richardson AE, Barea JM, McNeill AM, Prigent-Combaret C. 2009. Acquisition of phosphorus and nitrogen in the rhizosphere and plant growth promotion by microorganisms. Plant and Soil 321: 305–339.

Riesenfeld CS, Schloss PD, Handelsman J. 2004. Metagenomics: Genomic Analysis of Microbial Communities. Annual Review of Genetics 38: 525–552.

Risely A. 2020. Applying the core microbiome to understand host–microbe systems. Journal of Animal Ecology 89: 1549–1558.

Rodríguez H, Fraga R, Gonzalez T, Bashan Y. 2006. Genetics of phosphate solubilization and its potential applications for improving plant growth-promoting bacteria. Plant and Soil 287: 15–21.

Rout GR, Sahoo S. 2015. Role of Iron in Plant Growth and Metabolism. Reviews in Agricultural Science 3: 1–24.

Rungin S, Indananda C, Suttiviriya P, Kruasuwan W, Jaemsaeng R, Thamchaipenet A. 2012. Plant growth enhancing effects by a siderophore-producing endophytic streptomycete isolated from a Thai jasmine rice plant (Oryza sativa L. cv. KDML105). Antonie van Leeuwenhoek 102: 463–472.

Salas-González I, Reyt G, Flis P, Custódio V, Gopaulchan D, Bakhoum N, Dew TP, Suresh K, Franke RB, Dangl JL, et al. 2021. Coordination between microbiota and root endodermis supports plant mineral nutrient homeostasis. Science 371: 151–156.

Santos-Medellín C, Edwards J, Liechty Z, Nguyen B, Sundaresan V. 2017. Drought Stress Results in a Compartment-Specific Restructuring of the Rice Root-Associated Microbiomes. mBio 8: 1–15.

Santos-Medellín C, Liechty Z, Edwards J, Nguyen B, Huang B, Weimer BC, Sundaresan V. 2021. Prolonged drought imparts lasting compositional changes to the rice root microbiome. Nature Plants.

Sasse J, Martinoia E, Northen T. 2018. Feed Your Friends: Do Plant Exudates Shape the Root Microbiome? Trends in Plant Science 23: 25–41.

Schmidt W, Thomine S, Buckhout TJ. 2020. Editorial: Iron Nutrition and Interactions in Plants. Frontiers in Plant Science 10: 1–4.

Schneijderberg M, Schmitz L, Cheng X, Polman S, Franken C, Geurts R, Bisseling T. 2018. A genetically and functionally diverse group of non-diazotrophic Bradyrhizobium spp. colonizes the root endophytic compartment of Arabidopsis thaliana. BMC Plant Biology 18: 61.

Schreiner RP, Scagel CF. 2017. Leaf Blade versus Petiole Nutrient Tests as Predictors of Nitrogen, Phosphorus, and Potassium Status of ‘Pinot Noir’ Grapevines.

Shade A, Stopnisek N. 2019. Abundance-occupancy distributions to prioritize plant core microbiome membership. Current Opinion in Microbiology 49: 50–58.

Simons M, Permentier HP, De Weger LA, Wijffelman CA, Lugtenberg BJJ. 1997. Amino acid synthesis is necessary for tomato root colonization by Pseudomonas fluorescens strain WCS365. Molecular Plant-Microbe Interactions 10: 102–106.

Singh D, Geat N, Rajawat MVS, Mahajan MM, Prasanna R, Singh S, Kaushik R, Singh RN, Kumar K, Saxena AK. 2018. Deciphering the Mechanisms of Endophyte-Mediated Biofortification of Fe and Zn in Wheat. Journal of Plant Growth Regulation 37: 174–182.

Sprent JI. 2007. Evolving ideas of legume evolution and diversity: a taxonomic perspective on the occurrence of nodulation. New Phytologist 174: 11–25.

Swift JF, Hall ME, Harris ZN, Kwasniewski MT, Miller AJ. 2021. Grapevine Microbiota Reflect Diversity among Compartments and Complex Interactions within and among Root and Shoot Systems. Microorganisms 9: 92.

Swift JF, Migicovsky Z, Trello GE, Miller AJ. 2023. Grapevine bacterial communities display compartment-specific dynamics over space and time within the Central Valley of California. Environmental Microbiome 18: 84.

Talwar C, Nagar S, Kumar R, Scaria J, Lal R, Negi RK. 2020. Defining the Environmental Adaptations of Genus Devosia: Insights into its Expansive Short Peptide Transport System and Positively Selected Genes. Scientific Reports 10: 1–18.

Unno Y, Shinano T. 2013. Metagenomic Analysis of the Rhizosphere Soil Microbiome with Respect to Phytic Acid Utilization. Microbes and Environments 28: 120–127.

Verdugo-Vásquez N, Gutiérrez-Gamboa G, Villalobos-Soublett E, Zurita-Silva A. 2021. Effects of Rootstocks on Blade Nutritional Content of Two Minority Grapevine Varieties Cultivated under Hyper-Arid Conditions in Northern Chile. Agronomy 11: 327.

Viaene T, Langendries S, Beirinckx S, Maes M, Goormachtig S. 2016. Streptomyces as a plant’s best friend? FEMS Microbiology Ecology 92: 1–10.

Vink SN, Dini-Andreote F, Höfle R, Kicherer A, Salles JF. 2021. Interactive effects of scion and rootstock genotypes on the root microbiome of grapevines (Vitis spp. l.). Applied Sciences (Switzerland) 11: 1–11.

Walters WA, Jin Z, Youngblut N, Wallace JG, Sutter J, Zhang W, González-Peña A, Peiffer J, Koren O, Shi Q, et al. 2018. Large-scale replicated field study of maize rhizosphere identifies heritable microbes. Proceedings of the National Academy of Sciences 115: 7368–7373.

Wang S, Guo Z, Wang L, Zhang Y, Jiang F, Wang X, Yin L, Liu B, Liu H, Wang H, et al. 2021. Wheat Rhizosphere Metagenome Reveals Newfound Potential Soil Zn-Mobilizing Bacteria Contributing to Cultivars’ Variation in Grain Zn Concentration. Frontiers in Microbiology 12: 1–13.

Wang N, Li H, Wang B, Ding J, Liu Y, Wei Y, Li J, Ding G-C. 2022. Taxonomic and Functional Diversity of Rhizosphere Microbiome Recruited From Compost Synergistically Determined by Plant Species and Compost. Frontiers in Microbiology 12: 1–12.

Warschefsky EJ, Klein LL, Frank MH, Chitwood DH, Londo JP, von Wettberg EJB, Miller AJ. 2016. Rootstocks: Diversity, Domestication, and Impacts on Shoot Phenotypes. Trends in Plant Science 21: 418–437.

Wasai-Hara S, Hara S, Morikawa T, Sugawara M, Takami H, Yoneda J, Tokunaga T, Minamisawa K. 2020. Diversity of Bradyrhizobium in Non-Leguminous Sorghum Plants: B. ottawaense Isolates Unique in Genes for N2O Reductase and Lack of the Type VI Secretion System. Microbes and Environments 35.

Wei T, Simko V. 2021. R package ‘corrplot’: Visualization of a Correlation Matrix.

Wickham H, Averick M, Bryan J, Chang W, McGowan LD, François R, Grolemund G, Hayes A, Henry L, Hester J, et al. 2019. Welcome to the Tidyverse. Journal of Open Source Software 4: 1686.

Wolpert JA, Smart DR, Anderson M. 2005. Lower Petiole Potassium Concentration at Bloom in Rootstocks with Vitis berlandieri Genetic Backgrounds. American Journal of Enology and Viticulture 56: 163–169.

Wolters B, Jacquiod S, Sørensen SJ, Widyasari-Mehta A, Bech TB, Kreuzig R, Smalla K. 2018. Bulk soil and maize rhizosphere resistance genes, mobile genetic elements and microbial communities are differently impacted by organic and inorganic fertilization. FEMS Microbiology Ecology 94: 1–13.

Wright AH, Ali S, Migicovsky Z, Douglas GM, Yurgel S, Bunbury-Blanchette A, Franklin J, Adams SJ, Walker AK. 2022. A Characterization of a Cool-Climate Organic Vineyard’s Microbiome. Phytobiomes Journal 6: 69–82.

Xu L, Dong Z, Chiniquy D, Pierroz G, Deng S, Gao C, Diamond S, Simmons T, Wipf HML, Caddell D, et al. 2021. Genome-resolved metagenomics reveals role of iron metabolism in drought-induced rhizosphere microbiome dynamics. Nature Communications 12: 3209.

Xu L, Naylor D, Dong Z, Simmons T, Pierroz G, Hixson KK, Kim Y-M, Zink EM, Engbrecht KM, Wang Y, et al. 2018a. Drought delays development of the sorghum root microbiome and enriches for monoderm bacteria. Proceedings of the National Academy of Sciences 115: E4284–E4293.

Xu J, Zhang Y, Zhang P, Trivedi P, Riera N, Wang Y, Liu X, Fan G, Tang J, Coletta-Filho HD, et al. 2018b. The structure and function of the global citrus rhizosphere microbiome. Nature Communications 9: 4894.

Yagi S, Kitai S, Kimura T. 1971. Oxidation of elemental sulfur to thiosulfate by Streptomyces. Applied microbiology 22: 157–159.

Yanagita T, Foster JW. 1956. A BACTERIAL RIBOFLAVIN HYDROLASE. Journal of Biological Chemistry 221: 593–607.

Yeoh YK, Dennis PG, Paungfoo-Lonhienne C, Weber L, Brackin R, Ragan MA, Schmidt S, Hugenholtz P. 2017. Evolutionary conservation of a core root microbiome across plant phyla along a tropical soil chronosequence. Nature Communications 8.

Yoon J-H, Kang S-J, Park S, Oh T-K. 2007. Devosia insulae sp. nov., isolated from soil, and emended description of the genus Devosia. International Journal of Systematic and Evolutionary Microbiology 57: 1310–1314.

Yu X, Cloutier S, Tambong JT, Bromfield ESP. 2014. Bradyrhizobium ottawaense sp. nov., a symbiotic nitrogen fixing bacterium from root nodules of soybeans in Canada. International Journal of Systematic and Evolutionary Microbiology 64: 3202–3207.

Yu S-M, Lo S-F, Ho T-HD. 2015. Source–Sink Communication: Regulated by Hormone, Nutrient, and Stress Cross-Signaling. Trends in Plant Science 20: 844–857.

Yurgel SN, Rice J, Domreis E, Lynch J, Sa N, Qamar Z, Rajamani S, Gao M, Roje S, Bauer WD. 2014. Sinorhizobium meliloti flavin secretion and bacteria-host interaction: Role of the bifunctional RibBA protein. Molecular Plant-Microbe Interactions 27: 437–445.

Zhalnina K, Louie KB, Hao Z, Mansoori N, da Rocha UN, Shi S, Cho H, Karaoz U, Loqué D, Bowen BP, et al. 2018. Dynamic root exudate chemistry and microbial substrate preferences drive patterns in rhizosphere microbial community assembly. Nature Microbiology 3: 470–480.

Zhong H, Liu Z, Zhang F, Zhou X, Sun X, Li Y, Liu W, Xiao H, Wang N, Lu H, et al. 2022. Metabolomic and transcriptomic analyses reveal the effects of self- and hetero-grafting on anthocyanin biosynthesis in grapevine. Horticulture Research 9: uhac103.

Ziegler G, Terauchi A, Becker A, Armstrong P, Hudson K, Baxter I. 2013. Ionomic Screening of Field-Grown Soybean Identifies Mutants with Altered Seed Elemental Composition. The Plant Genome 6.

