## Supplemental Appendix for "Conserved Rhizosphere Microbiomes and Metabolic Functions Across Diverse Grapevine Rootstocks: Implications for Plant Elemental Composition"

**This PDF file includes:**

Supplementary Figures S1 to S10

Tables S1 to S7

### **Supplementary Figures**

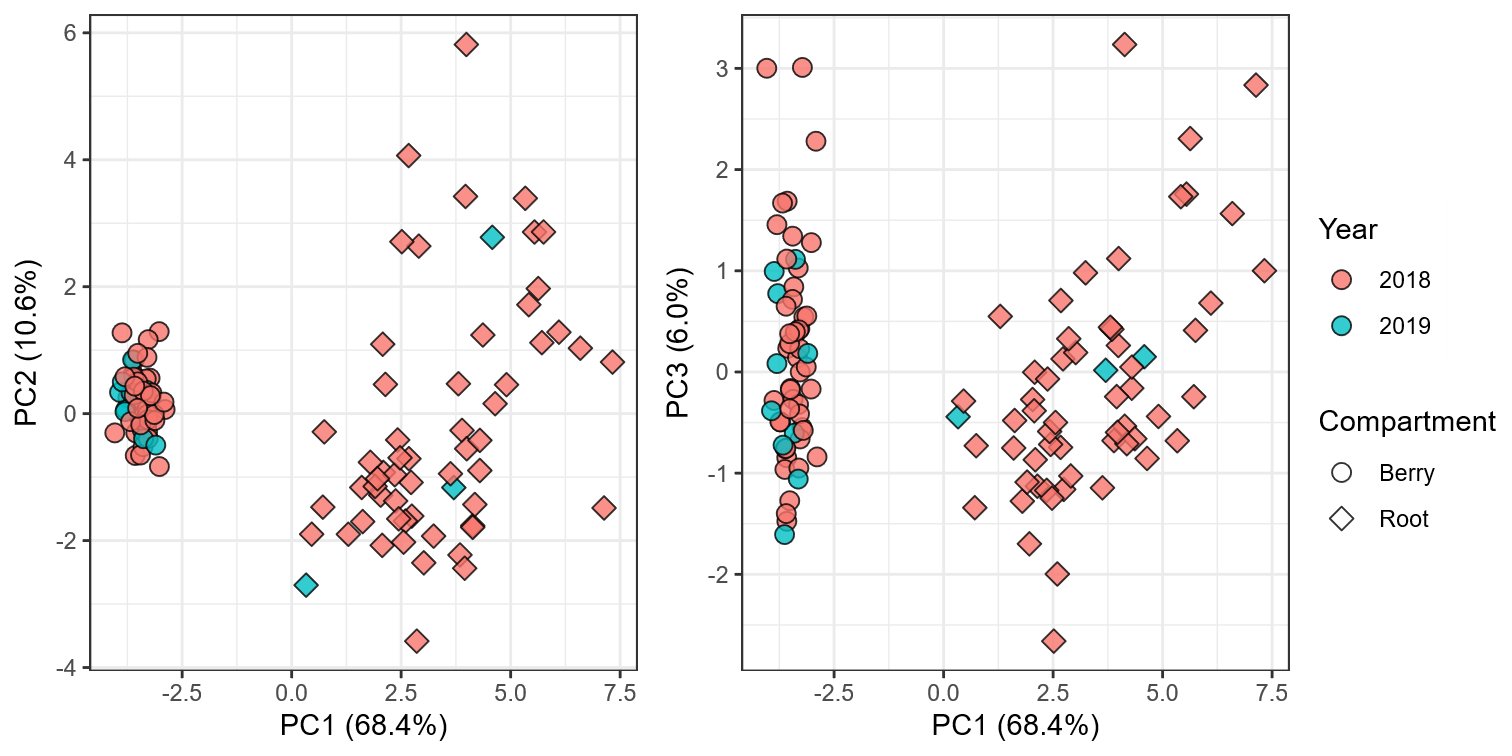

**Figure S1. The year of collection does not show clustering for either plant compartment.** Principal component analysis for elemental composition. Points are colored by the year they were collected, and plant compartments are represented by shapes.

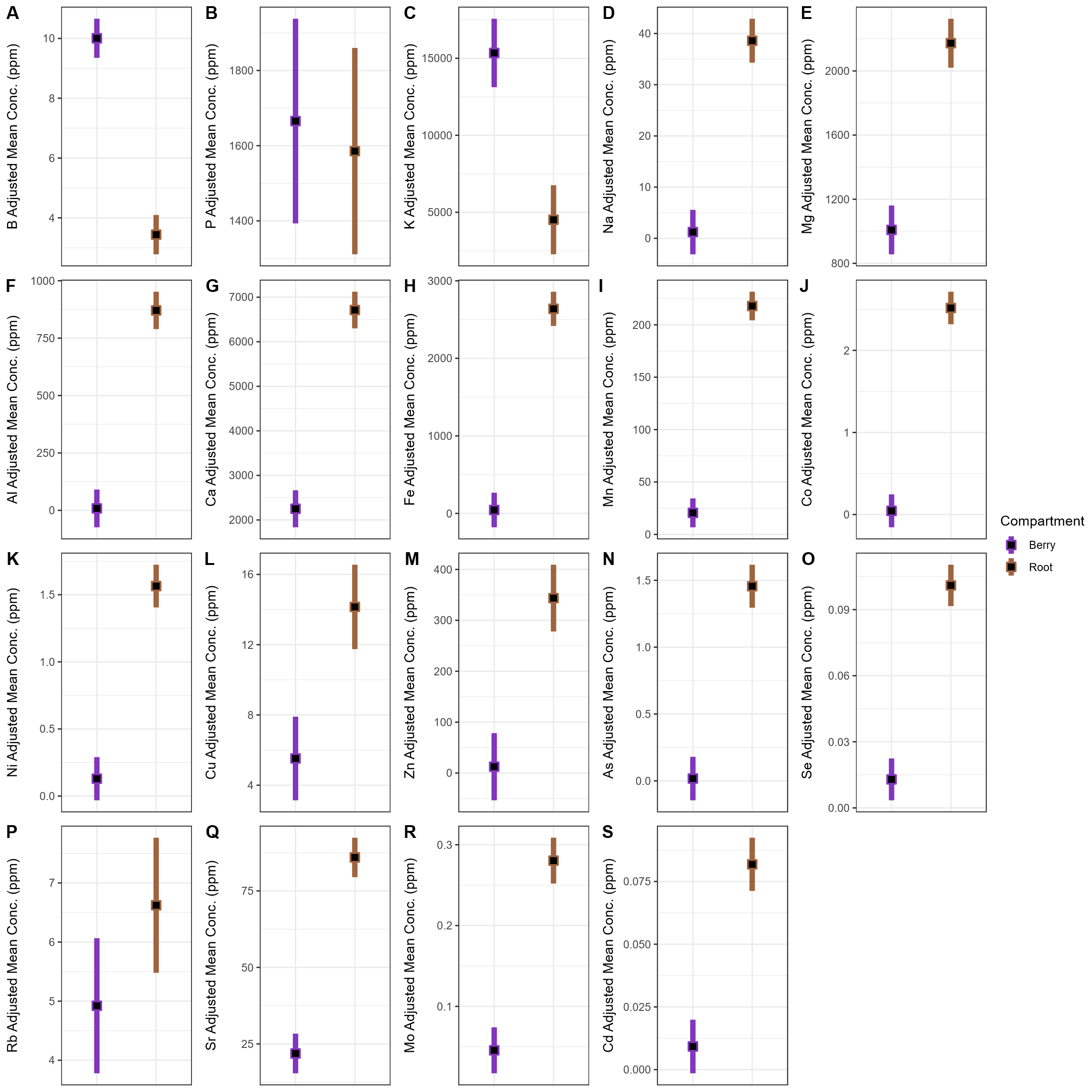

**Figure S2. Elemental concentrations by compartment for all elements.** Black squares represent estimated marginal means (EMM) for each compartment and colored bars show the 95% confidence intervals for the EMMs.

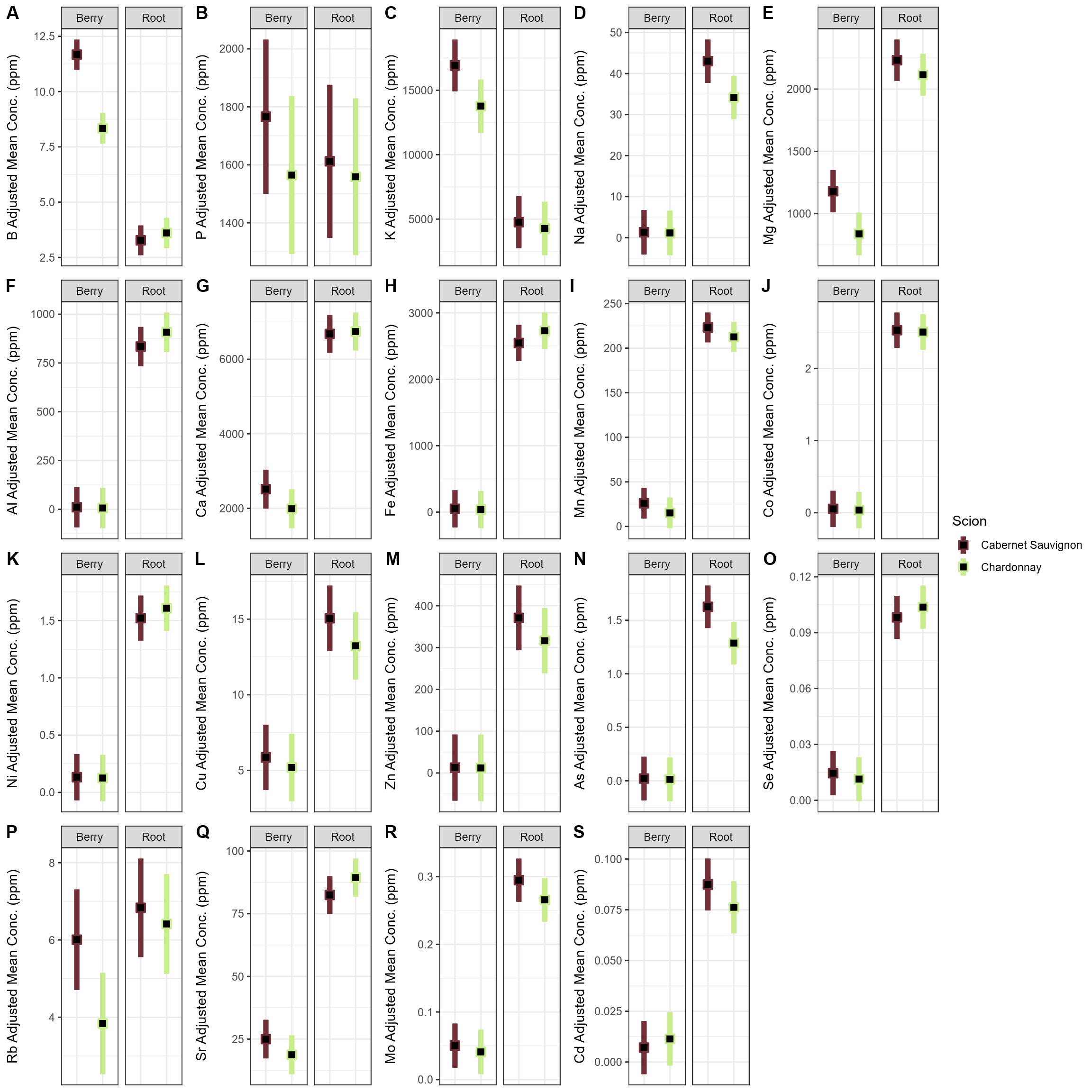

**Figure S3. Elemental concentrations by scion and compartment for all elements.** Black squares represent estimated marginal means (EMM) for each compartment and colored bars show the 95% confidence intervals for the EMMs.

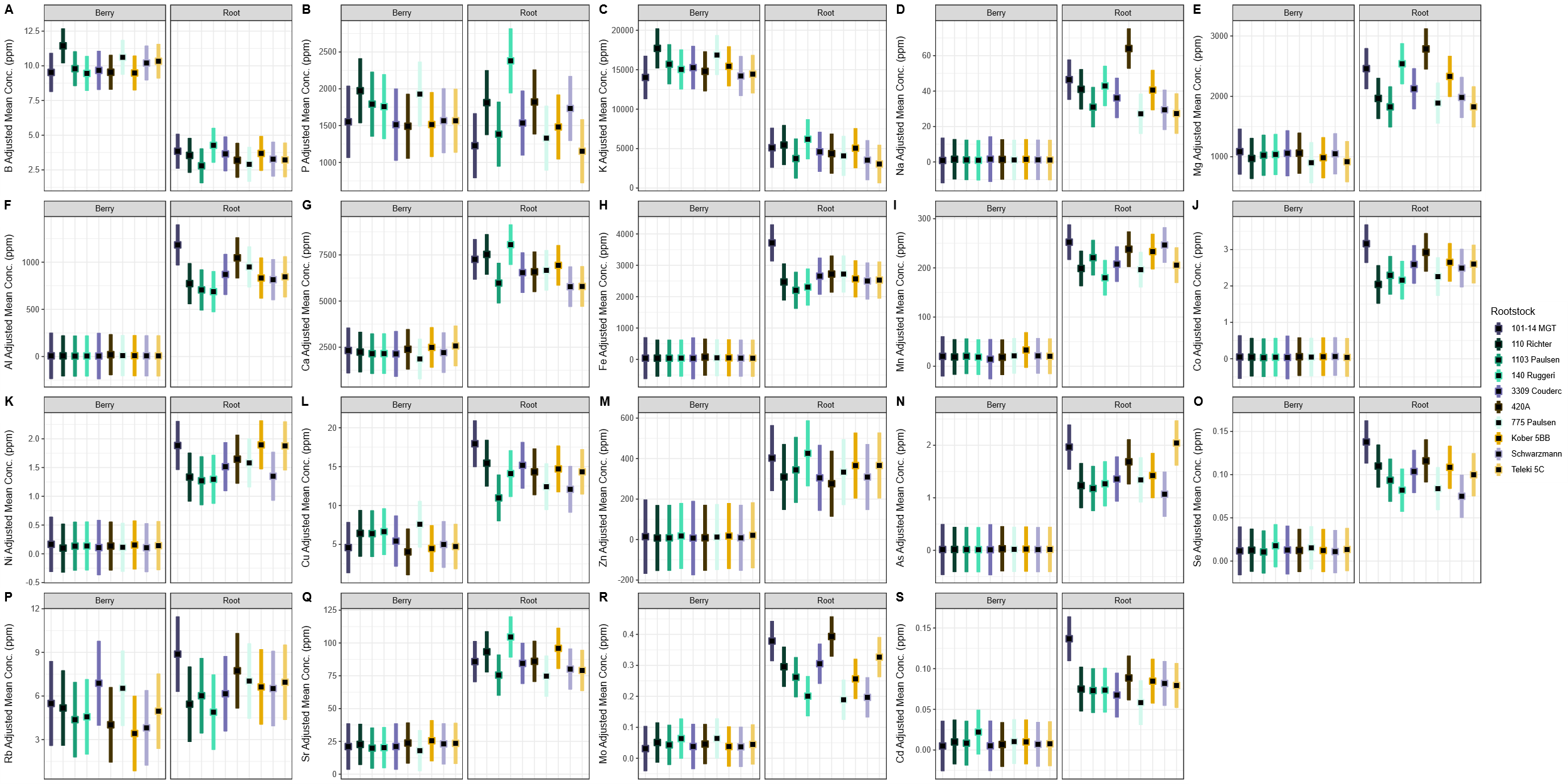

**Figure S4. Elemental concentrations by rootstock and compartment for all elements.** Black squares represent estimated marginal means (EMM) for each compartment and colored bars show the 95% confidence intervals for the EMMs.

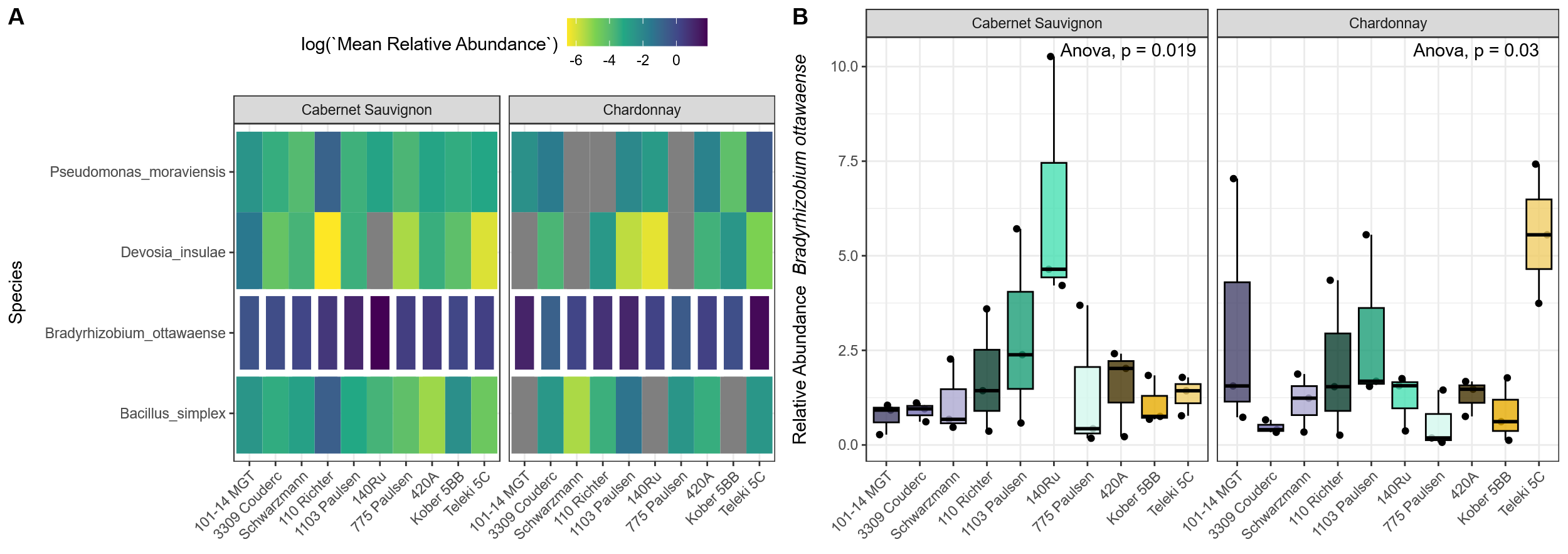

**Figure S5. Differentially abundance of species across rootstock and scion genotypes. A)** a heatmap depicts the mean relative abundance of species with significant differential abundance across the host genotypes. White borders highlight *Bradyrhizobium ottawaense* and **B)** a boxplot shows this taxa’s individual sample relative abundance.

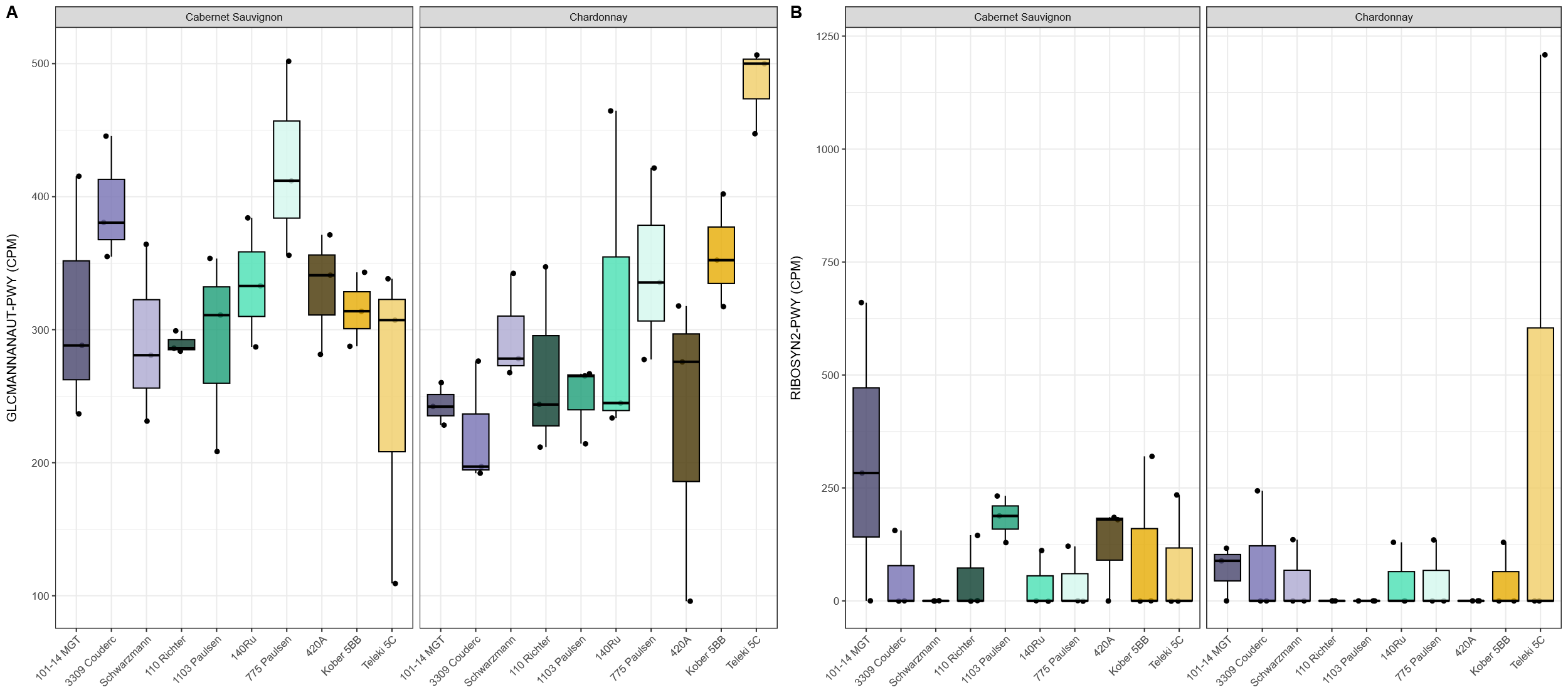

**Figure S6. Metabolic pathways exhibiting differential enrichment across rootstock/scion combinations.** A) superpathway of N-acetylglucosamine, N-acetylmannosamine and N-acetylneuraminate degradation (BioCyc ID: GLCMANNANAUT-PWY; ANOVA; *“Rootstock×Scion” p* < 0.01, F_9,40_ = 3.63) and B) superpathway of pyrimidine deoxyribonucleotides de novo biosynthesis (BioCyc ID: PWY-7211; ANOVA; *“Rootstock×Scion” p* = 0.01, F_9,40_ = 2.88).

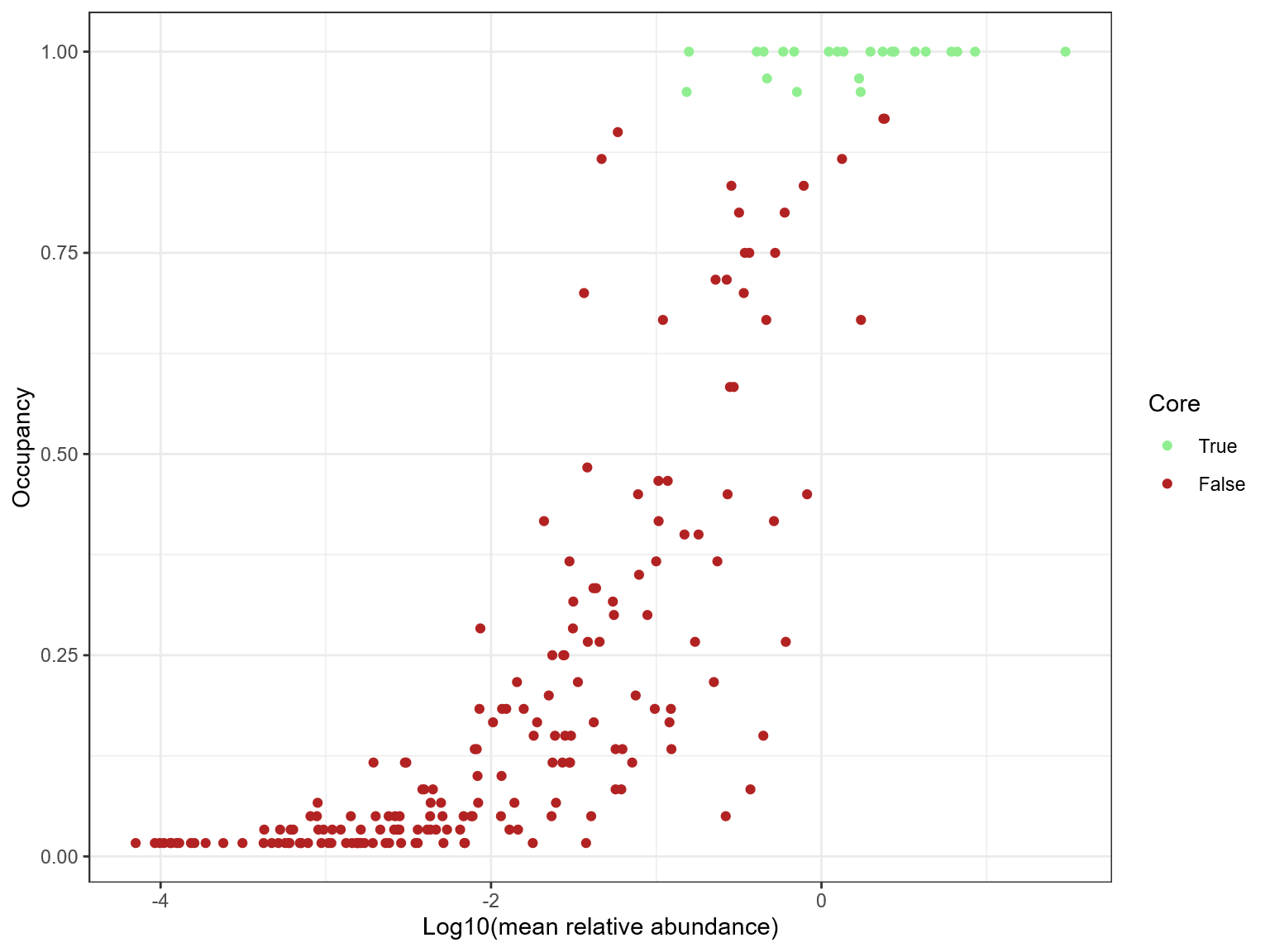

Figure S7. Abundance-occupancy distribution of genera in the rhizosphere microbiome. Core genera are defined as having an occupancy greater than 95% (i.e. 57/60 samples) and a mean relative abundance of 0.4% (log_10_ = -0.4).

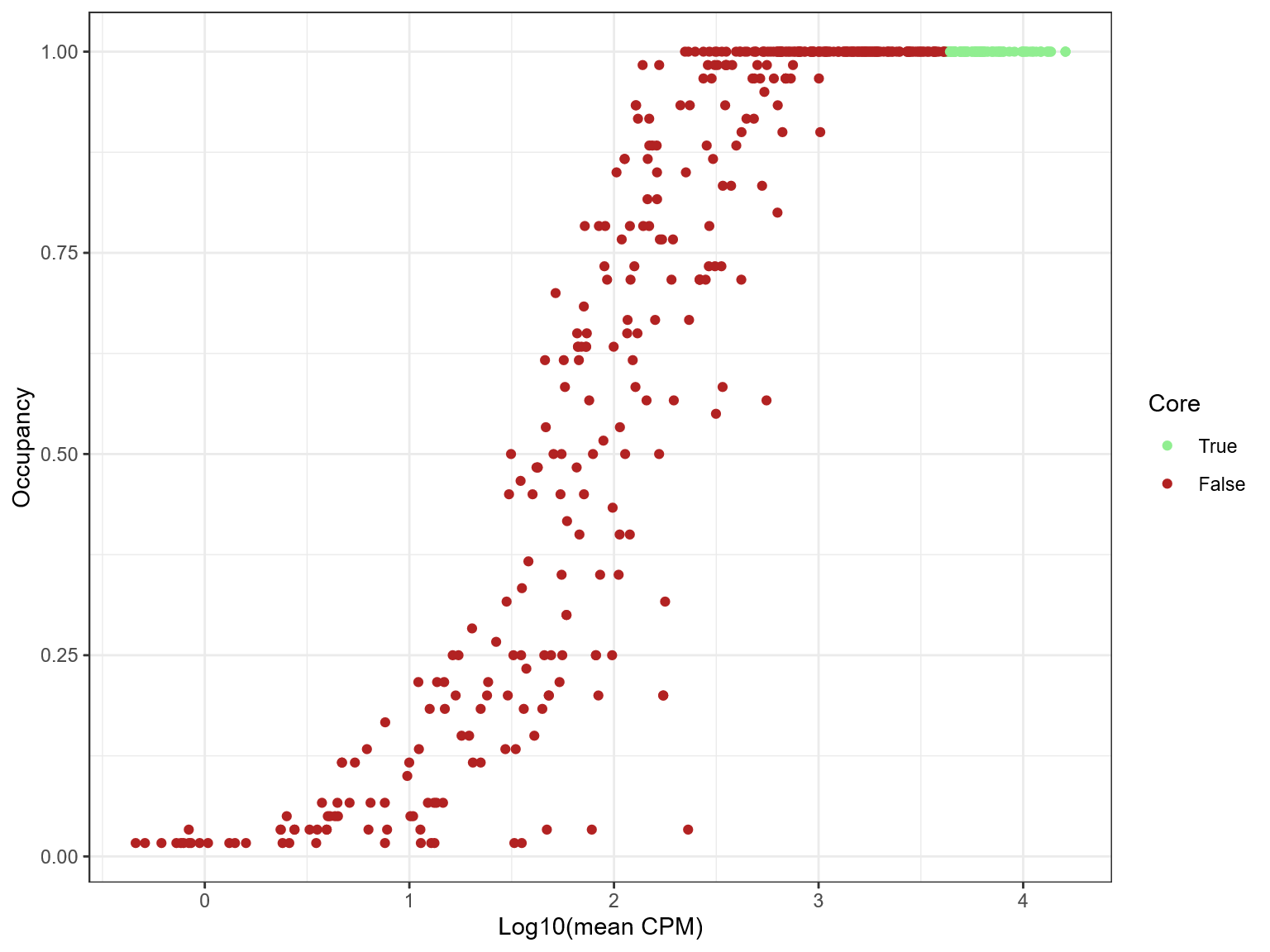

**Figure S8. Abundance-occupancy distribution of metabolic pathways in the rhizosphere microbiome.** Core pathways are defined as having an occupancy of 100% (*i.e.* 60/60 samples) and a mean CPM of 4215 (log_10_ = 3.62).

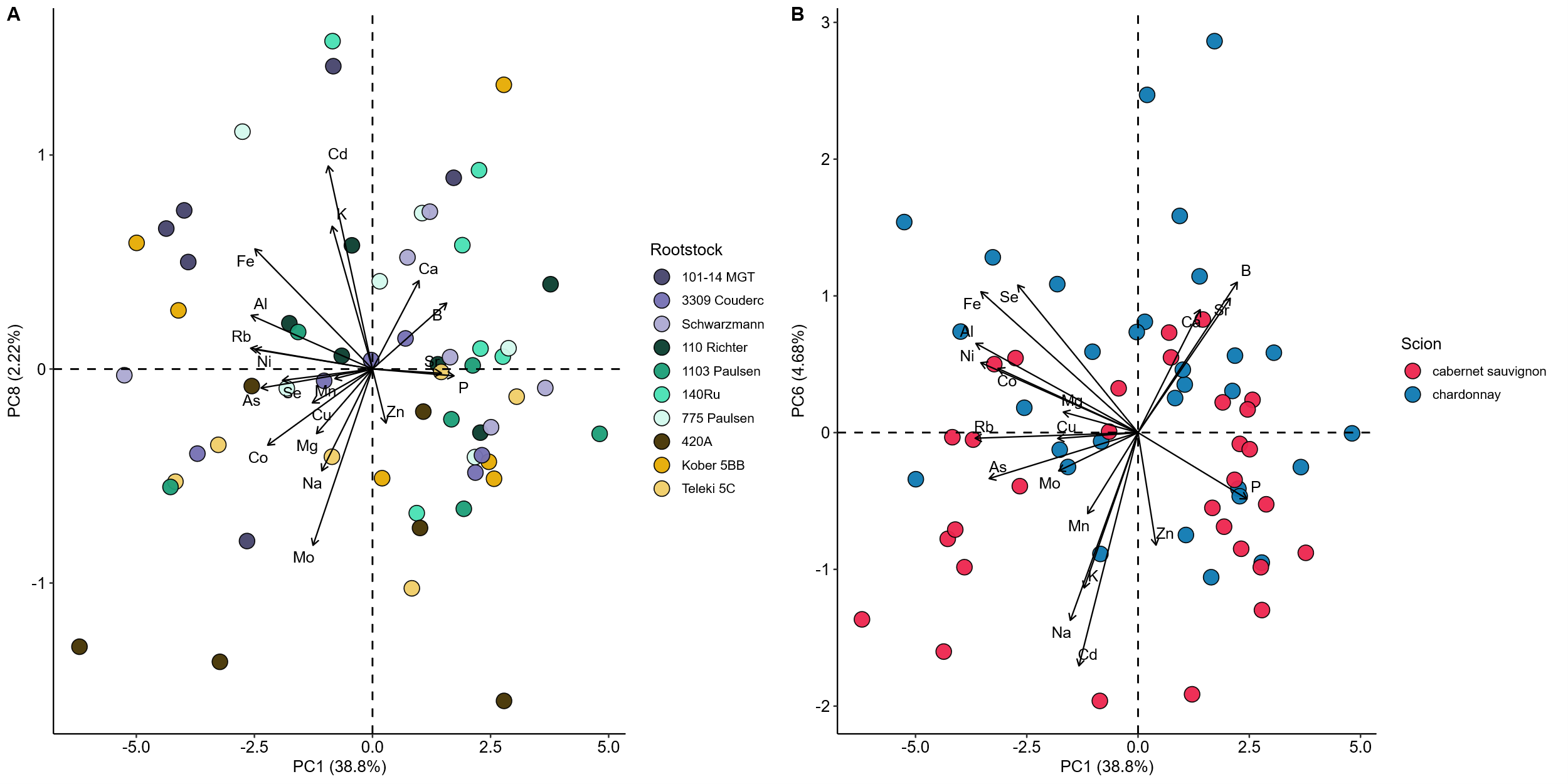

**Figure S9. Elemental composition principal components influenced by rootstock and scion genotype.** Each of the top ten PCs was treated as the response in a model with rootstock, scion genotype, and their interaction. A) PC8 was significantly influenced by rootstock genotype (ANOVA; *“Rootstock”* *p* < 0.01, F_9,22_ = 4.31). B) PC6 was significantly influenced by scion genotype (ANOVA; *“Scion”* *p* < 0.01, F_1,40_ = 10.71). PC1 is shown as the opposite axis in both plots. Arrows indicate loadings for each element.

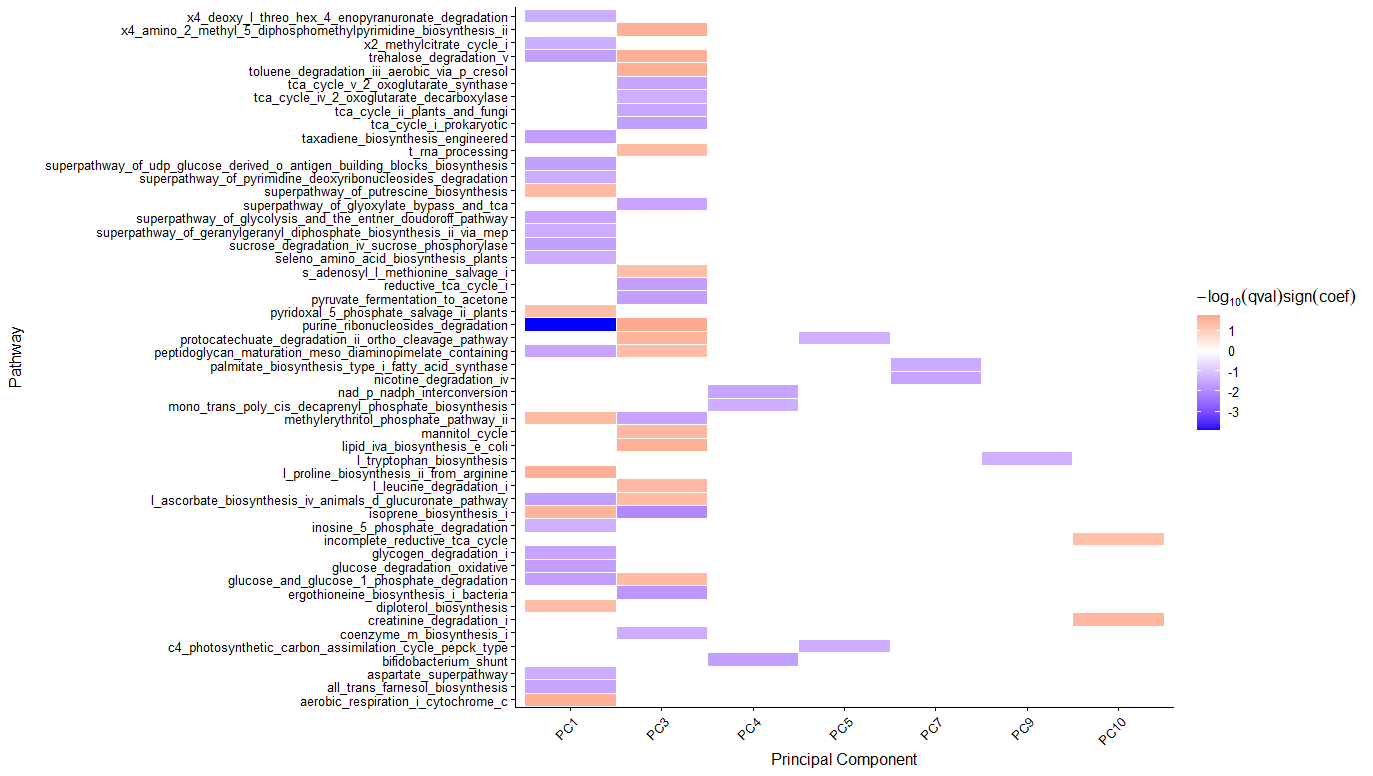

**Figure S10. Metabolic pathways associated with principal components.** Each of the top ten PCs, which were not associated with rootstock or scion genotype (see figure S9), was treated as the fixed effect in MaasLin2.

### **Supplementary Tables**

**Table S1. Parentage of rootstocks within the study.** Each is a cross between two of three *Vitis* spp.; *V. berlandieri* [Planch.], *V. riparia* [Michx]., and *V.* *rupestris* [Scheele]. Parentage information was compiled from the Vitis International Variety Catalogue (http://www.vivc.de/).

| **Rootstock** | **Parent 1** | **Parent 2** |
| --- | --- | --- |
| Kober 5BB | *V. berlandieri** | *V. riparia* |
| Teleki 5C | *V. berlandieri** | *V. riparia* |
| 420A | *V. berlandieri** | *V. riparia* |
| 101-14 MGT | *V. riparia* | *V. rupestris* |
| 3309 Couderc | *V. riparia* | *V. rupestris* |
| Schwarzmann | *V. riparia* | *V. rupestris* |
| 1103 Paulsen | *V. berlandieri** | *V. rupestris* |
| 110 Richter | *V. berlandieri** | *V. rupestris* |
| 140 Ruggeri | *V. berlandieri** | *V. rupestris* |
| 775 Paulsen | *V. berlandieri** | *V. rupestris* |

**Vitis berlandieri* is not formally recognized as a species and is synonymous with *Vitis cinerea* var *helleri* [(L.H. Bailey) M.O. Moore].

**Table S2. Random effect tables for element models.**

EXCEL FILE - No Table in Text

**Table S3. Type III ANOVA tables for element models.**

EXCEL FILE - No Table in Text

**Table S4. Type III ANOVA tables for element models with rootstock parentage groups.**

EXCEL FILE - No Table in Text

**Table S5.** Permutational multivariate ANOVA (PERMANOVA) using Bray-Curtis dissimilarity for rhizosphere microbiomes.

|  | **DF** | **SS** | **R^2^** | ***P*-value** |
| --- | --- | --- | --- | --- |
| **Rootstock** | 9 | 0.431 | 0.170 | 0.231 |
| **Scion** | 1 | 0.042 | 0.017 | 0.437 |
| **Residual** | 49 | 2.055 | 0.813 |  |

**Table S6. Core genera of the grapevine rhizosphere.** Genera are considered part of the core grapevine rhizosphere microbiome if they had an occupancy of 95% (*i.e.,* present in 57 of 60 samples) and a mean relative abundance of greater than 0.4% (80th percentile; See Figure S7).

| **Genera** | **Mean per sample relative abundance** | **Occupancy** |
| --- | --- | --- |
| *Rhizobium* | 30.00 | 1.00 |
| *Sphingomonas* | 8.51 | 1.00 |
| *Bacillus* | 6.64 | 1.00 |
| *Bradyrhizobium* | 6.11 | 1.00 |
| *Variovorax* | 4.28 | 1.00 |
| *Arthrobacter* | 3.68 | 1.00 |
| *Mesorhizobium* | 2.76 | 1.00 |
| *Streptomyces* | 2.66 | 1.00 |
| *Rhodococcus* | 2.35 | 1.00 |
| *Neorhizobium* | 1.98 | 1.00 |
| *Mycobacterium* | 1.73 | 0.95 |
| *Cupriavidus* | 1.69 | 0.97 |
| *Thermoleophilum* | 1.36 | 1.00 |
| *Inquilinus* | 1.25 | 1.00 |
| *Rhodospirillales* unclassified | 1.11 | 1.00 |
| *Pseudarthrobacter* | 0.71 | 0.95 |
| *Phyllobacterium* | 0.68 | 1.00 |
| *Paludisphaera* | 0.59 | 1.00 |
| *Rhodoplanes* | 0.47 | 0.97 |
| *Patulibacter* | 0.45 | 1.00 |
| *Kouleothrix* | 0.41 | 1.00 |

Table S7. Core metabolic pathways of the grapevine rhizosphere. Metabolic pathways are considered part of the core functional potential of grapevine rhizospheres if they had an occupancy of 100% and a mean enrichment of greater than 4215 CPM (80^th^ percentile; See Figure S8). Pathway enrichment values are given in copies per million (CPM), which accounts for gene length and number of mapped reads.

| **Pathway** | **Mean per sample CPM** |
| --- | --- |
| L-isoleucine biosynthesis I (from threonine) | 16095.723 |
| pyruvate fermentation to isobutanol (engineered) | 16095.723 |
| L-valine biosynthesis | 16095.723 |
| superpathway of branched chain amino acid biosynthesis | 13678.04 |
| L-isoleucine biosynthesis III | 13567.902 |
| adenosine deoxyribonucleotides de novo biosynthesis II | 13258.418 |
| guanosine deoxyribonucleotides de novo biosynthesis II | 13258.418 |
| superpathway of guanosine nucleotides de novo biosynthesis I | 13082.532 |
| guanosine ribonucleotides de novo biosynthesis | 12232.833 |
| superpathway of pyrimidine nucleobases salvage | 12184.309 |
| gondoate biosynthesis (anaerobic) | 12179.327 |
| adenosine ribonucleotides de novo biosynthesis | 11594.188 |
| superpathway of adenosine nucleotides de novo biosynthesis I | 11210.31 |
| superpathway of adenosine nucleotides de novo biosynthesis II | 11105.38 |
| superpathway of guanosine nucleotides de novo biosynthesis II | 10829.083 |
| fatty acid elongation -- saturated | 10478.698 |
| oleate biosynthesis IV (anaerobic) | 10319.114 |
| (5Z)-dodecenoate biosynthesis I | 10222.448 |
| cis-vaccenate biosynthesis | 9988.005 |
| 5-aminoimidazole ribonucleotide biosynthesis II | 9973.41 |
| superpathway of 5-aminoimidazole ribonucleotide biosynthesis | 9973.41 |
| palmitoleate biosynthesis I (from (5Z)-dodec-5-enoate) | 9823.229 |
| TCA cycle I (prokaryotic) | 9067.332 |
| folate transformations II (plants) | 8561.277 |
| superpathway of purine nucleotides de novo biosynthesis I | 8023.689 |
| superpathway of L-isoleucine biosynthesis I | 8019.014 |
| stearate biosynthesis II (bacteria and plants) | 7930.914 |
| superpathway of L-serine and glycine biosynthesis I | 7742.767 |
| L-arginine biosynthesis IV (archaebacteria) | 7711.227 |
| L-arginine biosynthesis I (via L-ornithine) | 7706.478 |
| L-arginine biosynthesis II (acetyl cycle) | 7578.822 |
| pyrimidine deoxyribonucleotides de novo biosynthesis IV | 7538.987 |
| pentose phosphate pathway (non-oxidative branch) I | 7413.81 |
| L-histidine biosynthesis | 7297.979 |
| folate transformations III (E. coli) | 7123.116 |
| UMP biosynthesis I | 7100.241 |
| superpathway of pyrimidine ribonucleotides de novo biosynthesis | 7031.327 |
| TCA cycle V (2-oxoglutarate synthase) | 7015.765 |
| tRNA charging | 6727.519 |
| pyrimidine deoxyribonucleotide phosphorylation | 6665.487 |
| inosine 5'-phosphate degradation | 6476.165 |
| inosine-5'-phosphate biosynthesis II | 6454.512 |
| superpathway of aromatic amino acid biosynthesis | 6439.458 |
| L-lysine biosynthesis III | 6403.541 |
| phosphopantothenate biosynthesis I | 6318.017 |
| octanoyl-[acyl-carrier protein] biosynthesis (mitochondria, yeast) | 6275.83 |
| 5-aminoimidazole ribonucleotide biosynthesis I | 6266.32 |
| inosine-5'-phosphate biosynthesis I | 6243.504 |
| L-lysine biosynthesis I | 6231.33 |
| L-ornithine biosynthesis I | 6228.376 |
| chorismate biosynthesis I | 6221.178 |
| superpathway of L-threonine biosynthesis | 6188.871 |
| TCA cycle IV (2-oxoglutarate decarboxylase) | 6184.169 |
| L-lysine biosynthesis VI | 6170.734 |
| chorismate biosynthesis from 3-dehydroquinate | 6065.66 |
| TCA cycle II (plants and fungi) | 5990.831 |
| superpathway of glyoxylate bypass and TCA | 5974.387 |
| glycolysis III (from glucose) | 5929.513 |
| glycolysis IV | 5835.749 |
| superpathway of L-lysine, L-threonine and L-methionine biosynthesis II | 5835.361 |
| adenine and adenosine salvage III | 5823.728 |
| superpathway of purine nucleotide salvage | 5801.142 |
| superpathway of acetyl-CoA biosynthesis | 5659.817 |
| assimilatory sulfate reduction I | 5635.604 |
| inosine-5'-phosphate biosynthesis III | 5591.375 |
| superpathway of glycolysis, pyruvate dehydrogenase, TCA, and glyoxylate bypass | 5365.902 |
| Rubisco shunt | 5320.377 |
| queuosine biosynthesis I (de novo) | 5306.339 |
| glycolysis I (from glucose 6-phosphate) | 5210.093 |
| superpathway of fatty acid biosynthesis initiation (E. coli) | 5204.975 |
| Calvin-Benson-Bassham cycle | 5179.41 |
| L-methionine biosynthesis III | 5153.725 |
| gluconeogenesis III | 5124.566 |
| glyoxylate cycle | 5085.218 |
| L-arginine biosynthesis III (via N-acetyl-L-citrulline) | 5006.693 |
| superpathway of coenzyme A biosynthesis III (mammals) | 4990.051 |
| superpathway of L-methionine biosynthesis (by sulfhydrylation) | 4955.613 |
| UDP-N-acetyl-D-glucosamine biosynthesis I | 4936.044 |
| tetrapyrrole biosynthesis II (from glycine) | 4910.443 |
| preQ0 biosynthesis | 4902.748 |
| peptidoglycan biosynthesis I (meso-diaminopimelate containing) | 4658.715 |
| UDP-N-acetylmuramoyl-pentapeptide biosynthesis I (meso-diaminopimelate containing) | 4646.172 |
| methylerythritol phosphate pathway I | 4640.079 |
| peptidoglycan biosynthesis III (mycobacteria) | 4615.815 |
| heme b biosynthesis I (aerobic) | 4608.89 |
| UDP-N-acetylmuramoyl-pentapeptide biosynthesis II (lysine-containing) | 4605.391 |
| glycolysis II (from fructose 6-phosphate) | 4592.531 |
| urea cycle | 4580.885 |
| tRNA processing | 4493.749 |
| L-citrulline biosynthesis | 4475.211 |
| superpathway of coenzyme A biosynthesis I (bacteria) | 4426.619 |
| gluconeogenesis I | 4399.371 |
| aerobic respiration I (cytochrome c) | 4388.6 |
| L-tryptophan biosynthesis | 4386.736 |
| anaerobic energy metabolism (invertebrates, cytosol) | 4381.354 |
| peptidoglycan maturation (meso-diaminopimelate containing) | 4278.444 |
| L-proline biosynthesis II (from arginine) | 4233.987 |
| C4 photosynthetic carbon assimilation cycle, PEPCK type | 4218.607 |
| dTDP-&beta;-L-rhamnose biosynthesis | 4215.218 |
